# Partners in space: Discordant population structure between legume hosts and rhizobium symbionts in their native range

**DOI:** 10.1101/2021.06.30.449460

**Authors:** Alex B. Riley, Michael A. Grillo, Brendan Epstein, Peter Tiffin, Katy D. Heath

## Abstract

Coevolution is predicted to depend on how the genetic diversity of interacting species is geographically structured. Plant-microbe symbioses such as the legume-rhizobium mutualism are ecologically and economically important, but distinct life history and dispersal mechanisms for these host and microbial partners, plus dynamic genome composition in bacteria, present challenges for understanding spatial genetic processes in these systems. Here we study the model rhizobium *Ensifer meliloti* using a hierarchically-structured sample of 191 strains from 21 sites in the native range and compare its population structure to that of its host plant *Medicago truncatula*. We find high local genomic variation and minimal isolation by distance across the rhizobium genome, particularly at the two symbiosis elements pSymA and pSymB, which have evolutionary histories and population structures that are similar to each other but distinct from both the chromosome and the host. While the chromosome displays weak isolation by distance, it is uncorrelated with hosts. Patterns of discordant population structure among elements with the bacterial genome has implications for bacterial adaptation to life in the soil versus symbiosis, while discordant population genetic structure of hosts and microbes might restrict local adaptation of species to each other and give rise to phenotypic mismatches in coevolutionary traits.

## Introduction

Coevolution is fundamentally a geographically variable process. This understanding has become widely accepted over the past several decades, most prominently with the inception of Thompson’s geographic mosaic theory of coevolution (Thompson, 1994, 2005). Coevolving partners can vary in their distributions, abundance, and genetic structure across the landscape and over time (Carlsson-Granér & Thrall, 2015; Fernandes, Lemos-Costa, Guimarães, Thompson, & de Aguiar, 2019; Laine, 2005; Tack, Horns, & Laine, 2014; Thompson, 2005). The evolution of interacting species will depend on spatial variation in selection, genetic variation, and gene flow. While there is abundant evidence for the core predictions of the GMTC (*i.e.,* selection mosaics, coevolutionary hot/cold spots, variable trait remixing) it is surprising that comparative analysis of population genetic structure among interacting partners has received relatively little attention (but see Caldera and Currie 2012; Baums et al. 2014; Strobel et al. 2016; Harrison et al. 2017).

Indeed, genetic structure can, in part, determine the outcomes of coevolution. For example, alignment in the geographic scale and location of population genetic structure between interacting species can allow for reciprocal selection and adaptation (*i.e.,* coevolutionary hotspots); conversely, discordant population structure, due to different levels of gene flow or other processes, can limit the ability of each species to locally adapt to the other and give rise to trait mismatches (Fernandes et al., 2019; Gandon, Capowiez, Dubois, Michalakis, & Olivieri, 1996a; Thompson, 2005). Thus, understanding the comparative population genetic structure of interacting species provides a key perspective on whether traits are expected to coevolve and how coevolutionary trait variation is maintained through time and space (Heath and Stinchcombe 2014; Hollowell et al. 2016a; Stoy et al. 2020).

Coevolutionary interactions typically occur between species with distinct life histories, range sizes, mating systems, and modes of dispersal □ all of which will impact population genetic structure (Pita, Rix, Slaby, Franke, & Hentschel, 2018; Revillini, Gehring, & Johnson, 2016; Thrall, Hochberg, Burdon, & Bever, 2007). In general, the majority of coevolutionary research has focused on systems involving antagonistic coevolution of eukaryotes. Mutualisms between macro and micro-organisms have received less attention, even though these mutualisms play critical roles in community and ecosystem processes (Berg, 2009; Revillini et al., 2016; Reynolds, Packer, Bever, & Clay, 2003).

The investigation of host-microbe interactions poses various logistical and methodological challenges. One challenge is the difficulty of isolating populations of a single species from the environment - particularly for soil bacteria in hyper-diverse communities (Heath & Grillo, 2016). In addition, many bacterial genomes are multipartite (Harrison et al. 2010; diCenzo and Finan 2017), including non-chromosomal replicons containing variable amounts of genetic information – ranging from small facultative plasmids that are often lost or transferred to large obligate segments containing core genes (often as megaplasmids or chromids; Harrison et al. 2010). Such divided genomes might represent an adaptation allowing functional division between replicons (reviewed by diCenzo and Finan 2017). Multipartite genome organization presents a challenge in the study of bacterial (co)evolution because different elements may move between individuals independent of each other, resulting in distinct evolutionary histories and population structure across different elements. These differences are compounded by the fact that these elements can also have widely differing rates of evolution and patterns of recombination leading to higher rates of evolution in plasmids not required for survival (Cooper, Vohr, Wrocklage, & Hatcher, 2010; Epstein et al., 2012; Epstein, Sadowsky, & Tiffin, 2014; Epstein & Tiffin, 2021).

The legume-rhizobium mutualism presents a tractable model system for studying microbial symbiosis due to the relative ease of isolating populations and genetic resources for several legume-rhizobium partnerships (Kaneko et al., 2002; Yates et al., 2015). In this symbiosis, rhizobia form nodules on plant roots, wherein they fix atmospheric N in exchange for carbon derived from plant photosynthesis; however, rhizobia inhabit the soil as saprophytes when not in symbiosis with a host, thereby experiencing profoundly different environmental conditions and likely selective pressures (Burghardt, 2020). In rhizobia with divided genomes (e.g., *Rhizobium* and *Ensifer* spp.), genes for symbiosis are carried on symbiosis elements (one or more megaplasmids or chromids), while the chromosome is thought to be responsible for core metabolic functions. These elements can have distinct evolutionary histories, diversity levels, and recombination rates (e.g., Bailly et al. 2011; Epstein et al. 2012, 2014; Cavassim et al. 2020; Epstein and Tiffin 2021).

Here we examine population structure in *Ensifer meliloti* (Becker et al., 2009; Tang et al., 2014), the rhizobium symbiont of the model legume *Medicago truncatula* (hereafter “*Medicago*”). *Medicago* is a self-fertilizing annual native to the Mediterranean region of Europe (Bonnin, Huguet, Gherardi, Prosperi, & Olivieri, 1996; Siol, Prosperi, Bonnin, & Ronfort, 2008). Research involving *Medicago* population genetics has identified a pattern of isolation by distance, and genetic structure at both the population and regional levels (Bonhomme et al., 2015; Bonnin et al., 1996; Grillo, De Mita, Burke, Solórzano-Lowell, & Heath, 2016; Ronfort et al., 2006; Siol et al., 2008). Here we use whole genome sequences for 191 isolates of *E. meliloti* sampled from 21 sites in the native range to characterize the population structure of this model mutualist and compare it to the host by reanalyzing *Medicago* sequence data from Grillo et al. (2016) to directly compare population structure in the two species. We ask: 1) how is genetic variation structured in *E. meliloti*; 2) do the three genomic elements of the rhizobium *E. meliloti* share common patterns of evolutionary history and population structure; and 3) do *Medicago* hosts and their rhizobium symbionts have correlated population genetic structure?

## Methods

### Study system

*Ensifer* (formerly *Sinorhizobium*) *meliloti* is a rhizobium species in the Alphaproteobacteria (Young & Haukka, 1996) that forms N-fixing nodules on the roots of multiple species in the genus *Medicago*, and is one of two *Ensifer* species that forms root nodules on *M. truncatula* (Zribi, Mhamdi, Huguet, & Aouani, 2004). The genome of *E. meliloti* is ∼6.79Mb, divided between three genomic elements: the chromosome (3.69Mb), megaplasmid pSymA (1.41Mb), and chromid pSymB (1.69Mb), as well as smaller plasmids in some strains (Galibert et al. 2001, Nelson et al. 2018). Metabolic modeling, along with genetic manipulation of the *E. meliloti* genome, have shown that gene content differs functionally between the three genomic elements (diCenzo, MacLean, Milunovic, Golding, & Finan, 2014; Galibert et al., 2001). The chromosome is predicted to carry primarily genes related to core metabolic function in soil, pSymA carries the majority of genes required for symbiotic N fixation, and pSymB carries primarily genes thought to be important for life in rhizosphere environments (diCenzo et al., 2014).

### Sample collection

We isolated *E. meliloti* strains from 21 sites in the native range of *Medicago* using a hierarchical sampling design with populations ranging from 1 to approximately 1350 km apart (Fig. S1; Table S1). Of the 21 sites, eight corresponded to the *Medicago* sites studied in Grillo *et al*. (2016). At each site we collected soil surrounding the top six inches of the root system from unearthed host plants. In order to generate a hierarchical sampling design, with individual strains ranging from 0-1350km apart (sites ranging from 1-1350km apart), we sampled multiple plants from some sites (see table S1). To avoid cross-contamination within each site, the sampling shovel was wiped clean of excess soil between samples and was pierced into the ground adjacent to a plant numerous times before sampling soil. Between sampling locations, the shovel was sterilized with dilute bleach. Soil samples were kept at 4□ prior to isolating cultures.

*Ensifer* strains were isolated or “trapped” in the laboratory from the field samples following standard protocols (Vincent 1970; Heath 2010). In brief, *Medicago* seeds were nicked with a razor blade, surface sterilized with 30% bleach, rinsed with sterile water, and imbibed in sterile water for approximately 30 minutes. Seeds were then directly sown into a given soil sample housed in a sterilized, fully self-contained Magenta box (see Brown et al. 2020 for details). Magenta boxes were randomly placed in a temperature-controlled grow room (23° C) under artificial light set to 12-h□ days. After four weeks, plants were harvested, and the soil was washed from the roots. Individual nodules were removed with forceps, surface sterilized by soaking in 30% bleach for 10 minutes, and then rinsed with sterilized water. Surface sterilized nodules were crushed with sterilized forceps and streaked on tryptone-yeast (TY) media plates. Plates were incubated at 30□ for 48 hours, sterilized glass stir rods were then used to streak samples were streaked on to TY plates which were again incubated at 30□. Individual colonies were then picked and grown in liquid TY media, and these pure cultures were stored in cryotubes in 50% TY 50% glycerol and stored at -80□. Given variation among host genotypes in rhizobium infection rates due to G x G interactions (Batstone, Dutton, Wang, Yang, & Frederickson, 2017; Heath & Tiffin, 2009), we used 24 *Medicago* host genotypes to trap rhizobia. Because both *E. meliloti* and *E. medicae* infect the roots of *Medicago* in the native range, we used a post-PCR restriction enzyme (RsaI) digestion of the 16S to assign strains to species (following Biondi et al. 2003). Ultimately, we isolated 191 strains of *E. meliloti* presented in the current study. We hereafter refer to *E. meliloti* as simply “*Ensifer”* or “symbiont”.

### Sequencing

We extracted DNA from cultures of *Ensifer* grown in liquid TY media using the Qiagen DNeasy kit (Hilden, Germany) and sent samples to the DOE Joint Genome Institute (JGI) for sequencing (Berkeley, CA, USA). JGI prepared a paired end sequencing library for each strain, and sequenced samples on an Illumina HiSeq-2500 1TB platform (101nt read length; Illumina, Inc., San Diego, CA, USA). Of the 199 strains submitted to JGI, we received high quality whole genome sequences for 166. We re-grew the remaining 33 strains from frozen cultures (as above), extracted DNA using the Zymo Quick-DNA kit for Fungi or Bacteria (Irvine, CA, USA). These samples were sequenced (2 × 150 or paired end 150 nt read length) on the Novaseq 6000 platform (Illumina, Inc, San Diego, CA, USA) by the Roy J. Carver biotechnology center at the University of Illinois at Urbana-Champaign (USA). We successfully recovered quality genome sequences from 25 of these 33 isolates, for a total of 191 *E. meliloti* strains sequenced.

### Genome assembly, annotation, and SNP calling

To ensure high quality SNP calling and genome assembly, we trimmed PCR adaptors and removed PCR duplicates and PhiX contamination using HT-Stream (github.com/s4hts/HTStream), followed by further adaptor removal, removal of bases with quality scores < 30 from the ends of reads, and removal of reads < 80 bp long with TrimGalore! (github.com/FelixKrueger/TrimGalore). For calling SNPs in core genes (genes present in > 80% of strains), we aligned reads to the *E. meliloti* reference genome USDA1106 using BWA with default settings (Li & Durbin, 2009). We then used Freebayes (Garrison & Marth, 2012) to identify haplotype variants, which we split into SNPs using VCFtools (Danecek et al., 2011). We found 491,277 variable SNPs with variant qualities above 20. We then filtered SNPs to retain only those with depth values between 20 and 230, minor allele frequencies ≥ 0.009 (*i.e.,* present in at least two strains), and that were present in at least 80% of individuals, 72,311 SNPs remained after applying these filter (chromosome: 34,689 SNPs, pSymA: 15,162 SNPs, pSymB: 22,460 SNPs; see Table S2). For variable gene content, we assembled genomes *de novo* using SPADES (Bankevich et al., 2012) with default parameters, followed by annotation using PROKKA (Seemann, 2014) and scanned these annotations for presence-absence variants using default settings in ROARY (Page et al., 2015).

### Phylogenetic and Statistical Analyses

To explore the population structure of the three genomic elements of *Ensifer,* we first used principal components analysis (PCA). We used the glPca function in the adegenet (Jombart & Ahmed, 2011) library in R on a random subsample of 15k SNPs (to equilibrate dataset size) for each element of the *Ensifer* genome to naively cluster individuals by genome-wide similarity. We then plotted the positions of individuals along the first three axes of variation (using ggplot2; Fig. S2) to visually compare how genetic variation is distributed across elements (Wickham, 2016). To quantify genetic differences we calculated individual and population-level Dxy (Nei 1972) using the stamppNeisD function from the package StAMPP (Pembleton, Cogan, & Forster, 2013) on all variants for each element. To test whether these patterns of genetic distance among *Ensifer* strains were congruent across the three genome elements, we used these matrices of individual-based distance metrics in pairwise Mantel tests comparing the three *Ensifer* genome elements (chromosome vs. pSymA, chromosome vs. pSymB. pSymA vs. pSymB). Mantel tests were implemented in the R package ade4 (Dray & Dufour, 2007). To explore geographic structure in the variable genome (*i.e.,* genes present in only some strains), we performed an additional genetic PCA, as above, using the matrix of *Ensifer* gene presence-absence variants (91,840 genes) and plotted the first three principal components for visual inspection.

For each of the three elements, we built phylogenies based on the same random subsample of 15k core genome SNPs used for PCA. We used the neighbor joining (nj) function in the R package ape (Paradis & Schliep, 2019) with 1000 bootstrap replicates and visualized trees in FigTree (v.1.4.4 ) (Rambaut, 2018). Based on these preliminary results, we treated each element independently for remaining analyses.

We used AMOVA (poppr.amova) with clone correction and 10,000 random permutations (Kamvar, Tabima, & Gr□unwald, 2014) to partition the genetic variance among individuals, sites, and regions, as well as to assess the statistical significance of these levels of spatial genetic structure. To test the hypothesis of isolation by distance, we calculated Pearson correlations between individual-level D_XY_ and geographic distances between sampling sites.

To compare the spatial genetic structure of rhizobia and host plants, we reanalyzed RAD-seq data from the 192 *Medicago* genotypes studied in Grillo *et al* (2016). We first called SNPs using Stacks with default parameters (Catchen, Hohenlohe, Bassham, Amores, & Cresko, 2013), then filtered the resulting variants using VCFtools (Danecek et al., 2011) to ensure that all variants were present in at least 80% of lines, had minor allele frequencies > 0.05, and were > 5kb apart, 10,814 SNPs remained after these filters. We performed AMOVA, calculated individual and population-based D_XY_, and tested for isolation by distance in *Medicago* as detailed above for *Ensifer*. For the subset of eight sites for which we had both hosts and symbionts, we used Mantel tests to ask whether the three matrices of pairwise population D_XY_ values from symbionts (chromosome, pSymA, and pSymB) were correlated with that of the host, to test for congruent population genetic structure between hosts and symbionts.

## Results

To address the extent of phylogenetic congruence between the chromosome, pSymA, and pSymB of *Ensifer*, we first used pairwise Mantel tests of whether the matrices of genetic distances were correlated. At the individual level, we found non-significant correlations between the chromosome and pSymA and between the chromosome and pSymB, while the two symbiosis elements (pSymA and pSymB) were more strongly correlated (left column, Fig. 1), indicating distinct evolutionary histories of the chromosome versus pSymA and pSymB. Neighbor joining trees also reveal distinct evolutionary histories among the three genome elements. The topology of the chromosomal tree showed five tightly clustered groups of individuals, with almost all strains from Corsica closely related and forming a distinct group (orange cluster; Fig. 2a) and strains from Spain somewhat interspersed with those from France mostly in the blue and yellow clusters. The diversity of strains found in mainland France (the best sampled region) included representatives from each major chromosomal lineage; indeed, pink and purple clusters were found only in mainland France (except strain 710A from Corsica), mostly on the western Mediterranean coast (Fig. S1; Table S1).

**Figure 1.**
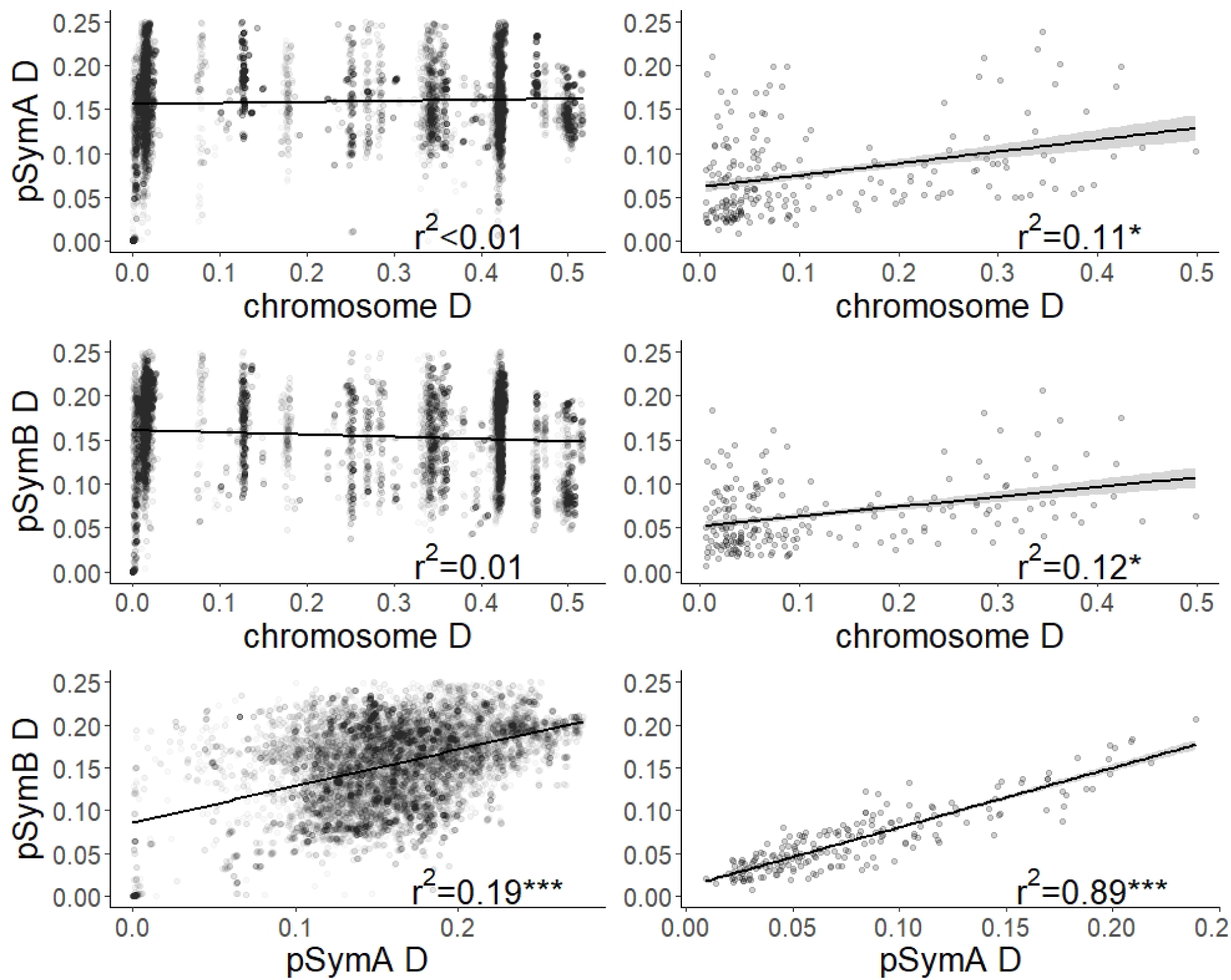
Tests of correlated population structure between the genomic elements in *E. meliloti*. Shown are correlations between the matrices of pairwise D_XY_ values. Left column are individual D_XY_ values and right column are population-level D_XY_ values.

**Figure 2.**
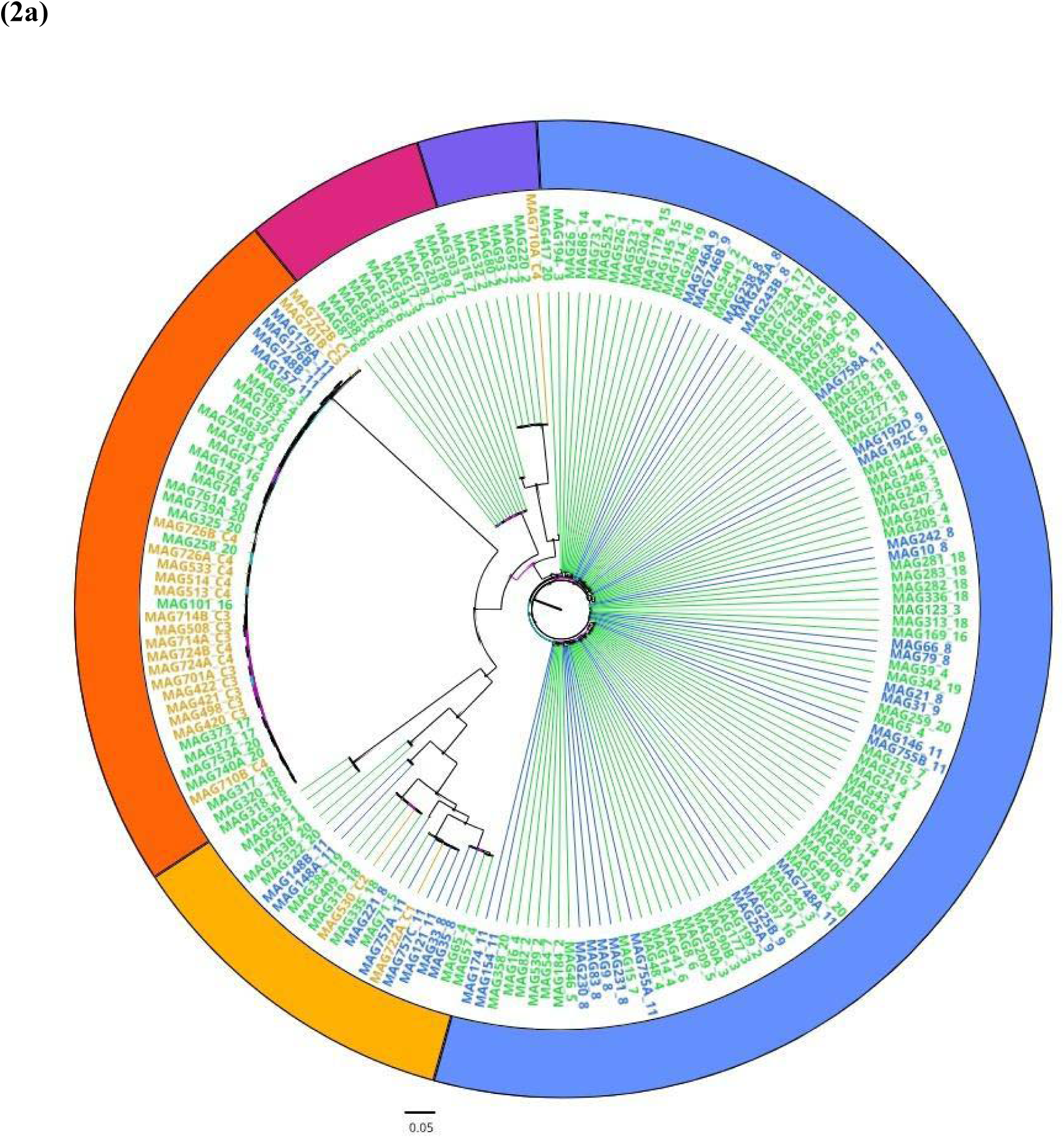
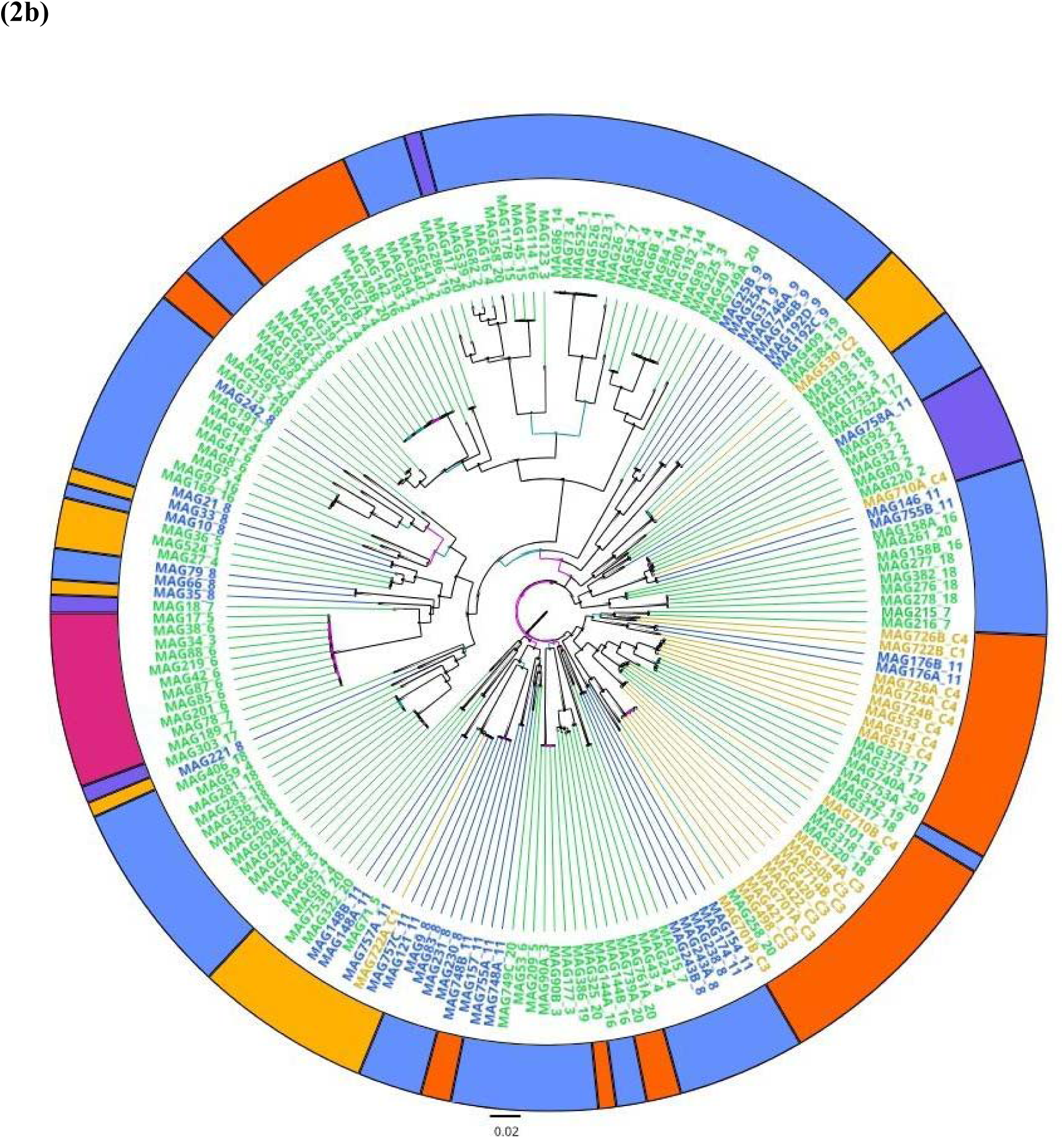
Neighbor joining trees of rhizobium individuals based on (a) chromosomal and (b) pSymA variant data. For both trees individual tip labels (strain ID followed by soil population) are colored based on region of origin (orange from Corsica, green from mainland France, blue from Spain). The outer ring coloration in both trees represents the five major clusters of the chromosome, allowing for comparison with pSymA. Only pSymA is shown here due to sharing a similar topology to pSymB (see Fig. S3).

Using these same colors (corresponding to the chromosomal clusters) to annotate the pSym trees (Fig. 2b; Fig. S3) facilitates visualization of differences in tree topology among the three elements, indicative of plasmid transfer (whole or partial) among chromosomal lineages. The internal branches of both symbiotic element trees reveal greater diversity, with chromosomal lineages spread across diverse pSym lineages in both trees (colored outer rings in Fig. 2b; Fig. S3). Some strains were closely related at all three elements of their genomes; for example, the group of closely related chromosomal lineages in pink from the western coast of France appear together in all three trees, indicating little pSym element recombination among these and other lineages (strain 78 in the pSymB tree is a notable exception). On the other hand, the other chromosomal clusters (purple, orange, and yellow clusters in Fig. 2a), appear throughout the pSymA and pSymB tree (see same colors interspersed; Fig. 2b; Fig. S3), indicating that these chromosomal lineages are found with diverse pSym genotypes. For example, the tightly clustered group of closely related chromosomal lineages in orange were found with lineages from across both the pSymA and pSymB trees.

Next, we used genetic PCA to explore population structure and AMOVA to partition the genome-wide genetic variation at the within-population, among-population, and among-region scales. Genetic PCA did not reveal strong differentiation in *Ensifer*, even at the regional scale, in either the core genome SNPs or variable gene content (Fig. S2; Fig. S4). Nevertheless, we found significant spatial genetic structure in the core genome at all scales investigated using AMOVA (Table 1), but that the patterns varied among the three *Ensifer* elements. For all three, we found considerable variation (66-76%) maintained within populations, compared to the among-population and among-region scales (Table 1). All three among-element correlations were stronger at the among-population level (right column, Fig. 1), revealing substantial congruence between pSymA and pSymB, and weakly correlated population structure between the chromosome and both symbiosis elements (right column, Fig. 1). The discrepancy between individual and population level correlations is likely due to high within population diversity (Table 1) and several populations with near-zero genetic distances (*e.g.,* within Spain and France, and between them Table S3).

**Table 1.**
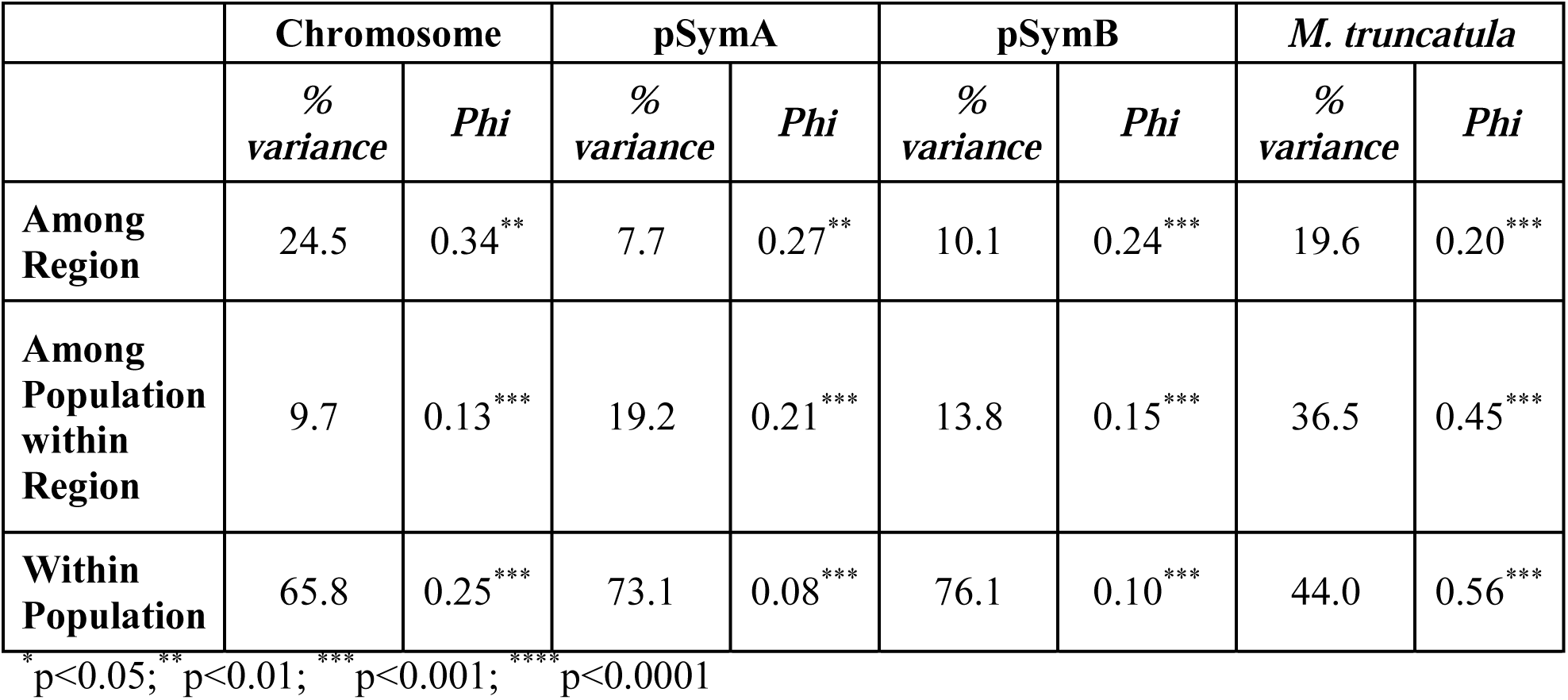
AMOVA partitioning the genome-wide genetic variation for each of the three elements of the *E. meliloti* genome for 191 strains sampled from 21 populations from three regions (Spain, France, or Corsica) in the native range of the symbiosis, compared to host plant *M. truncatula*. For each level of spatial division, the percent variance explained, phi statistic, and significance are given.

|  | Chromosome |  | pSymA |  | pSymB |  | *M. truncatula* |  |
| --- | --- | --- | --- | --- | --- | --- | --- | --- |
|  | %<br>variance | *Phi* | %<br>variance | *Phi* | %<br>variance | *Phi* | %<br>variance | *Phi* |
| **Among Region** | 24.5 | 0.34** | 7.7 | 0.27** | 10.1 | 0.24*** | 19.6 | 0.20*** |
| **Among Population within Region** | 9.7 | 0.13*** | 19.2 | 0.21*** | 13.8 | 0.15*** | 36.5 | 0.45*** |
| **Within Population** | 65.8 | 0.25*** | 73.1 | 0.08*** | 76.1 | 0.10*** | 44.0 | 0.56*** |

\* p<0.05; \*\* p<0.01; \*\*\* p<0.001; \*\*\*\* p<0.0001

The *Ensifer* chromosome showed the strongest differentiation at the regional scale (*i.e.,* among Spain, France, and Corsica; Table 1). This finding is likely driven by Corsica, as short branch lengths were found among most Corsica strains on the chromosomal tree (within the orange cluster; Fig 2a) and genetic distances were high between Corsica populations and the rest of the range (Table S3). The chromosome was also the only *Ensifer* element to exhibit significant isolation by distance (IBD) at this spatial scale (Fig 3). Despite overall IBD, there was extremely large variation in the genetic distances between *Ensifer* isolated from even the most distant locations (*i.e.,* D_XY_ distances ranged from 0-0.5 even when strains were found approximately 1350 km apart; Table 1). The two symbiotic elements in *Ensifer* (pSymA and pSymB) were less structured at the among-region and among-population scales (Table 1), and neither exhibited significant IBD at our sampling scale (Fig. 3). Thus although populations differed in genetic composition (significant structure in Table 1), *Ensifer* strains from distant populations up to 1350 km apart were often as closely related to each other as strains from the same sampling site.

**Figure 3.**
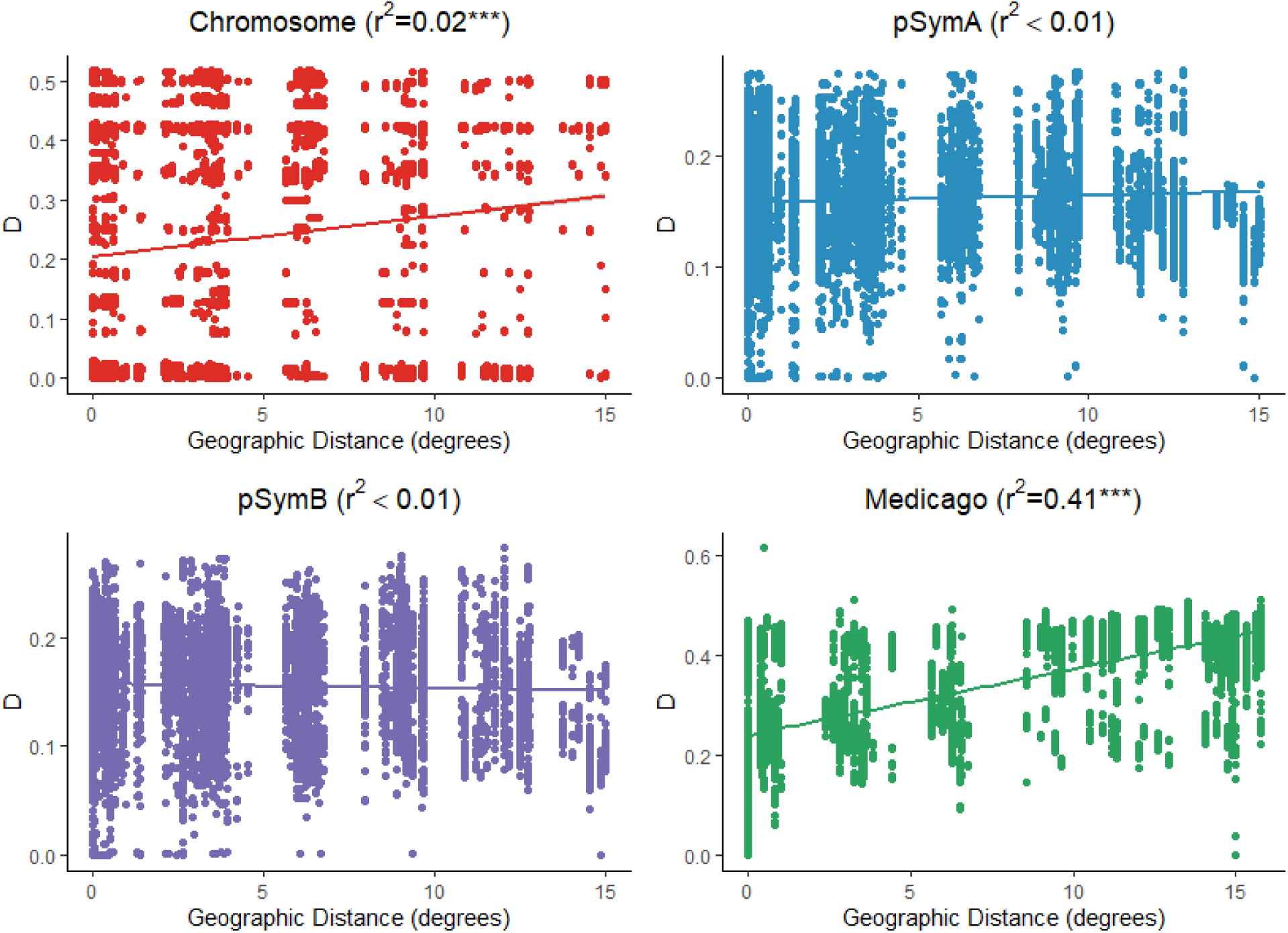
Correlations between geographic distance and individual genetic distance (D_XY_) for *Medicago* host plants (n=192) (green), and the *E. meliloti* (n=191) chromosome (red), pSymA (blue), and pSymB (purple). Shown are trend lines and Pearson r^2^ values from Mantel tests for each comparison (***p<0.001).

Hosts and symbionts had mismatched population genetic structure. We found no significant correlations between the pattern of among-population differentiation (D_XY_) between *Medicago* hosts and any element in the *Ensifer* genome (Fig. S5). While both the host and *Ensifer* chromosome exhibit patterns of isolation by distance (see above), their spatial patterns of genetic variation were distinct; while host plants primarily clustered along axes that separated Spain from France and Corsica (Fig. S2; Grillo et al. 2016 and references therein), the *Ensifer* chromosome most clearly clustered into two groups (mainland Europe versus Corsica along PC1, Fig. S2). The lack of IBD and little regional structure in pSymA and pSymB indicate that host plants at a given site have the potential to interact with the diversity of symbiosis plasmids from across this portion of the species range.

## Discussion

Evolution, and thus coevolution, is shaped by the interaction of selection with demographic processes such as gene flow occurring across a species range; therefore, understanding the maintenance of genetic variation, the evolution of mutualistic traits, and host-microbe interactions requires that we consider individuals and species in their spatial context. Here we used a hierarchically-structured sample of legume hosts and rhizobial symbionts to show: 1) the elements of the bacterial genome (chromosome versus pSymA and pSymB) have distinct population genetic structure at the local and regional scales, 2) host plants have more spatial genetic structure than rhizobia, particularly when compared to the two symbiotic elements (pSymA and pSymB), and 3) patterns of population structure between hosts and symbionts are not correlated across this portion of their native range.

### Spatial genetic processes at the bacterial chromosome and symbiotic elements

The population structure and dispersal abilities of microbes remain poorly understood (Chase et al., 2019; Green, Bohannan, & Whitaker, 2008; Hanson, Fuhrman, Horner-Devine, & Martiny, 2012; Martiny et al., 2006; VanInsberghe, Arevalo, Chien, & Polz, 2020). Sequence data have revealed considerable variation in dispersal ability among microbial taxa (Locey & Lennon, 2016). Based on metagenomics datasets, rhizobia have many characteristics associated with large geographic ranges and, by extension, large dispersal ability (*e.g.,* Proteobacteria, plant-associated or soil-borne taxa, and large genome sizes; Choudoir et al. 2018). However, metagenomic studies of microbial communities are limited in their resolution; therefore, population genomic data (Hoetzinger, Pitt, Huemer, & Hahn, 2021), coupled with spatially-structured samples of both within- and among-population variation (Whitaker & Banfield, 2006), are required for inferring population structure and gene flow, which are key to local adaptation and coevolution (Thompson 2005; Hoeksema and Forde 2008; Kraemer and Boynton 2017). Like other diverse microbial systems (Chase et al., 2019; Hoetzinger et al., 2021; Vos & Velicer, 2008), *Ensifer* displays both isolation by distance (chromosome) and population genetic structure (all elements); nevertheless, we found extremely closely-related strains up to 1350 km apart – indicating at least some long-distance dispersal. Large dispersal ability, together with the observation of abundant local phenotypic (Heath, 2010; Heath & Tiffin, 2009) and genomic (Bailly et al., 2011; Bailly, Olivieri, De Mita, Cleyet-Marel, & Béna, 2006) variation for symbiosis, suggests that most of the variation relevant to both bacterial adaptation and coevolution can be found at relatively small spatial scales.

At the regional scale sampled here, we identify distinct evolutionary histories and population genetic structures between the chromosome and the symbiosis elements, but similar population structure between the two symbiosis elements. Previous work has shown divergent evolutionary histories of the three elements in this species (Bailly et al. 2006; Galardini et al. 2013; Epstein et al. 2014; Nelson et al. 2018), as well as in other rhizobium taxa with separate symbiosis plasmids (Cavassim et al., 2020; Klinger, Lau, & Heath, 2016; Koppell & Parker, 2012; Kumar et al., 2015; Pérez Carrascal et al., 2016; Young et al., 2006) or symbiosis gene regions (Amanda C. Hollowell et al., 2016; Porter, Faber-Hammond, Montoya, Friesen, & Sackos, 2019). Our analyses support these findings while also revealing how the three elements differ at the within-population, among-population, and regional scale using a hierarchically-structured spatial sample of within and among-site variation. Using this approach allowed us to resolve distinct fine-scale spatial genetic processes at individual elements, namely geographically limited chromosomal lineages (e.g., Corsica), but little population structure of symbiosis elements.

One hypothesis for the existence of distinct elements in bacteria is to break up genetic correlations among traits that are important during different phases of the life cycle and/or establish linkage between adaptive gene complexes (diCenzo & Finan, 2017). Based on functional genetics and metabolic models (Galibert et al. 2003; diCenzo et al. 2014; diCenzo and Finan 2017), the three major genome elements are thought to play somewhat distinct roles in the life history of *Ensifer:* chromosome (basic metabolism), pSymA (nodulation and N fixation), and pSymB (rhizosphere). The traits, and potentially genes, that underlie fitness in the soil, rhizosphere, and nodules are likely distinct (Friesen and Mathias 2010; Sachs et al. 2011; Heath and Stinchcombe 2014; Burghardt 2019); having these processes segregated into distinct genome elements might allow the buildup of coadapted gene complexes within elements while facilitating independent adaptation to host and abiotic conditions on separate elements. One particularly interesting geographic disjunction that distinguishes the chromosome from the other elements is a cluster of extremely closely-related lineages that was nearly ubiquitous in Corsica but found elsewhere in low frequency. Although it remains unclear how this pattern arose, it is possible that this chromosomal lineage has swept to high frequency as the result of recent selection. Epstein et al. (2012) found low diversity along half of the *E. meliloti* chromosome in a range-wide sample of 24 strains, likely indicating a recent selective sweep with little recombination. Given little recombination on the chromosome (Epstein et al. 2012; Nelson et al. 2018), targets of selection on this element can be difficult to distinguish. Nevertheless it would be interesting to test whether the three genome elements of *Ensifer* are locally adapted in ways consistent with their functional gene composition, possibly by using host genetic variation alongside abiotic climatic factors in a landscape genomics model (e.g., Yoder et al. 2014; Rudman et al. 2018).

Our work suggests that pSymA and pSymB might be inherited together, leading to strong correlations between them at both the individual and population scales. Recombination in *Ensifer* is known to occur through conjugation of pSymA and pSymB (Ding & Hynes, 2009). Recent work indicates that pSymB is mobilized by pSymA via a *rctA*-mediated control system, though tracking these events in real time has proven elusive in the laboratory (Blanca-Ordóñez et al., 2010; diCenzo & Finan, 2017; Galibert et al., 2001; Pérez-Mendoza et al., 2005; Pretorius-Guth, Puhler, & Simon, 1990). If pSymA and pSymB are coupled in nature, this could give rise to genetic constraint that could affect *Ensifer*’s ability to adapt to distinct selective forces arising from various environmental conditions encountered in the rhizosphere versus in association with host plants. Functional predictions (*e.g.,* gene knockout experiments, functional annotation based on homology) do not always predict quantitative trait variation; therefore, hypotheses regarding genetic constraint among the elements should be addressed with quantitative genetic correlations between rhizosphere and symbiosis traits (e.g., Ossler and Heath 2018; Wood et al. 2018).

Distinct spatial genetic processes between the chromosome and symbiosis elements, and the coupling of the symbiosis elements, begs the question of how horizontal transmission of the symbiosis elements and recombination generate population structure in these bacteria. While we summarize the geographical patterns at the whole-element scale in order to compare these to the host (see below), finer-scaled work on gene tree heterogeneity (Degnan & Rosenberg, 2009) within each of the three elements is required for a full understanding of whether individual loci, large genomic regions, or even entire plasmids are introgressed through conjugation, and at what spatial scales such events occur. For example, Porter et al. (2019) recently used fine-scale patterns of gene co-occurrence across *Mesorhizobium* strains from California to discover the local transfer and duplication of symbiosis island and associate this variation with symbiotic partner quality.

### Mismatched host-symbiont population structure

Our results indicate discordance in population structure between hosts and symbionts due to differences in drift, gene flow, mutation, and/or spatially-variable selection. *Medicago* host plants are most structured along a longitudinal gradient, indicative of distinct glacial refugia, dividing populations in Spain from France and Corsica (Branca et al., 2011; Grillo et al., 2016; Ronfort et al., 2006; Yoder et al., 2014). By contrast, we found that most genome-wide variation in *Ensifer* is found within individual sites, and our phylogenetic and spatial analyses indicate that regional differentiation and IBD (which was significant for the chromosome) are different from host plants, since chromosomes from Corsica were distinct from most mainland populations (see discussion above). In contrast to host plants and the *Ensifer* chromosome, pSymA and pSymB showed substantially less regional differentiation and no pattern of IBD, due to increased introgression following long-range dispersal events, higher effective population size, or both.

A striking feature of mutualisms is the existence of abundant genetic variation for mutualistic traits within and among populations, given models that predict the erosion of trait variation in mutualism (Heath and Stinchcombe 2014; Stoy et al. 2020). Indeed considerable genetic variation in partner choice, sanctioning, signaling, and partner quality has been identified in the *Medicago-Ensifer* mutualism (Batstone et al., 2017; Batstone, O’Brien, Harrison, & Frederickson, 2020; Burghardt et al., 2018; Burghardt, Epstein, & Tiffin, 2019; Heath, 2010; Heath & Tiffin, 2009). A mismatch in the population genetic structure of interacting species, as we identified here, could restrict populations of host and symbionts from locally adapting to each other and thereby contribute to the maintenance of coevolutionary trait variation. Thus strong local selection would be necessary to overcome the homogenizing effects of gene flow (*i.e.,* migration-selection balance; Savolainen et al. 2013) in order for hosts and symbionts to locally adapt to each other. Empirical work in other systems comparing the population structure of interacting species finds a range of outcomes, from correlated spatial genetic patterns between partners (Anderson, Olivieri, Lourmas, & Stewart, 2004; Caldera & Currie, 2012; Smith, Godsoe, Tank, Yoder, & Pellmyr, 2008; Thompson, Thacker, & Shaw, 2005), to largely discordant patterns, where one species exhibits substantially less population structure than its partner (Baums et al., 2014; Dybdahl & Lively, 1996; Strobel et al., 2016).

Spatially-explicit theory on mutualism coevolution indicates that relative rates of gene flow between hosts and symbionts alters the likelihood of trait matching/local coadaptation, and even that gene flow can help explain the existence of maladaptation and abundant genetic variation for mutualism (Nuismer, Thompson, & Gomulkiewicz, 2003; Parker, 1999; Yoder & Nuismer, 2010). An important finding of these models, however, is that the outcomes of coevolution depend on how gene flow interacts with forms of natural selection, yet mutualism research often lacks explicit information on how selection in the wild acts on mutualism traits like partner quality and partner choice (Heath and Stinchcombe 2014; Stoy et al. 2020).

The degree of fitness alignment in mutualism, and thus the degree to which these interactions coevolve like better-studied antagonisms, is an unresolved question (Batstone et al., 2020; Frederickson, 2017; Maren L. Friesen, 2012; Gano-Cohen et al., 2020; Jones et al., 2015; Sachs, Quides, & Wendlandt, 2018). For antagonistic interactions, which are governed by negative frequency-dependent selection due to interspecific conflict, models suggest that the partner with a higher migration rate will benefit from the influx of new, potentially adaptive alleles that provide a competitive edge (Carlsson-Granér & Thrall, 2015; Gandon, Capowiez, Dubois, Michalakis, & Olivieri, 1996b). If conflict prevails in mutualisms, and increases in rhizobium fitness occur at the expense of the host (Gano-Cohen et al., 2020; Porter & Simms, 2014), incongruent population structures between hosts and symbionts should make it harder for hosts to evolve mechanisms to exclude lower-quality symbionts (*e.g.,* partner choice; Akçay 2017; Younginger and Friesen 2019), though this depends on the strength of selection. On the other hand, alignment between host and symbiont fitness might be common in mutualisms, as suggested by recent lab and experimental evolution studies (Batstone et al., 2020; Friesen, 2012). This scenario should ultimately result in positive frequency-dependent selection wherein novelty is disfavored (unless populations are far from their adaptive peak), leading to stable allele frequencies under purifying selection. Mismatched population structure, as identified here, would restrict the efficacy of positive frequency-dependence and any resulting local adaptation between hosts and symbionts. Our finding of discordant population genetic structure is in agreement with other host-microbe symbioses where microbes are transmitted horizontally and independently of their hosts (Dybdahl and Lively 1996; Baums et al. 2014; Strobel et al. 2016; Harrison et al. 2017). If this is common, then coevolutionary selection would have to be quite strong to overcome the gene flow to generate patterns of local coadaptation.

## Acknowledgments

We thank Laurène Gay and Joelle Ronfort for advice and contributions to field collection, and Amy Marshall-Colón, Julian Catchen, and Rachel Whitaker for feedback on an early manuscript draft. We thank various sources of funding including NSF IOS-1401864 to MAG, NSF IOS-1645875 to KDH, NSF PGRP-1856744 to PT and KDH, JGI CSP-503446 to PT and KDH, and the Department of Plant Biology at the University of Illinois Urbana-Champaign.

## Data accessibility

Raw sequence reads and assemblies are archived at NCBI (accessions ###).

## Author contributions

KDH and MAG conceived of and designed the work. KDH, MAG, and PT acquired funding. MAG collected soil, performed trapping experiments, and isolated strains, and MAG and ABR extracted and submitted DNA for sequencing. ABR and BE performed bioinformatic and evolutionary analyses. ABR, KDH, and MAG drafted the article, and all authors participated in critical revisions and approved the final version for submission.

---

**Table S1.**
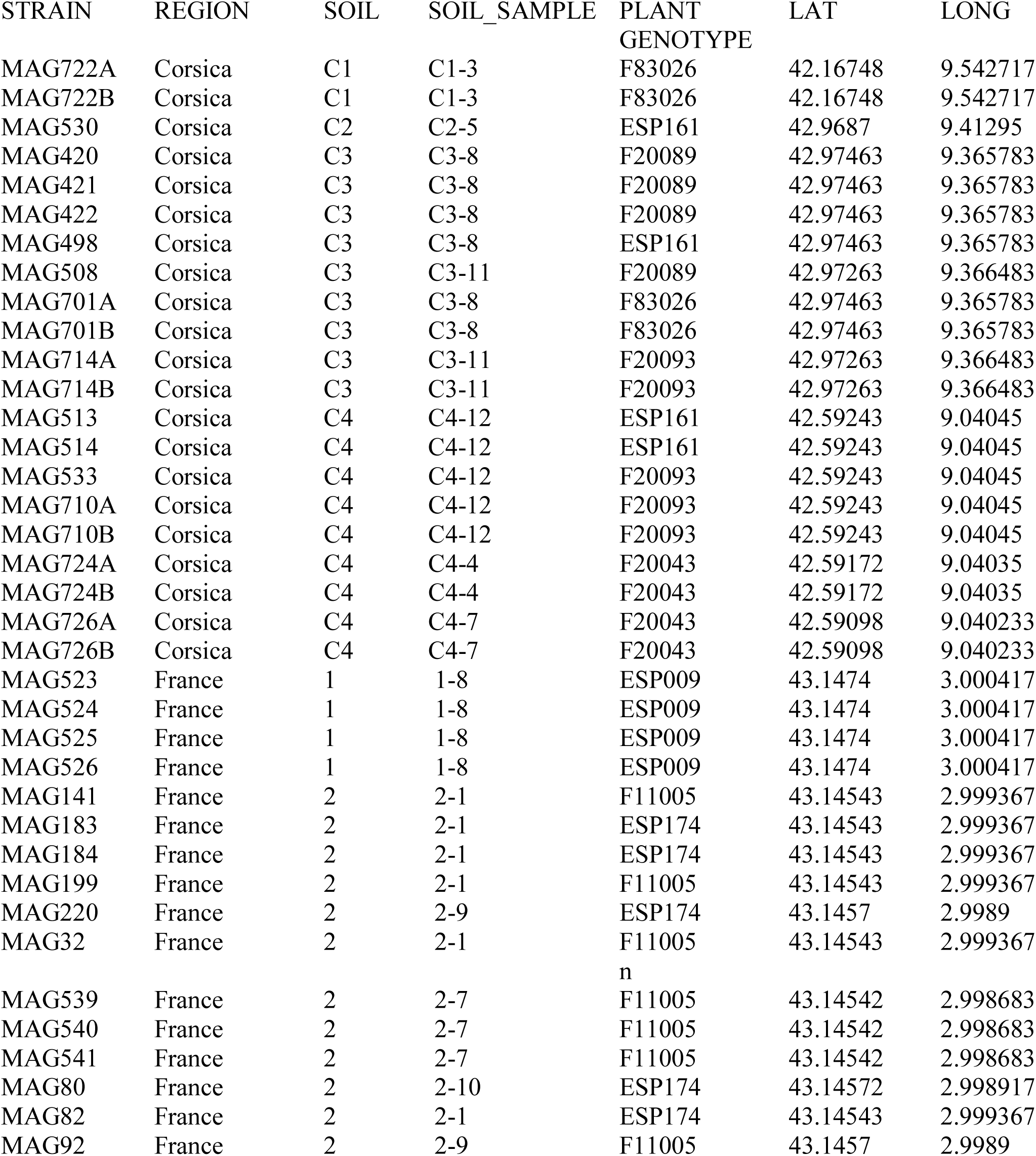
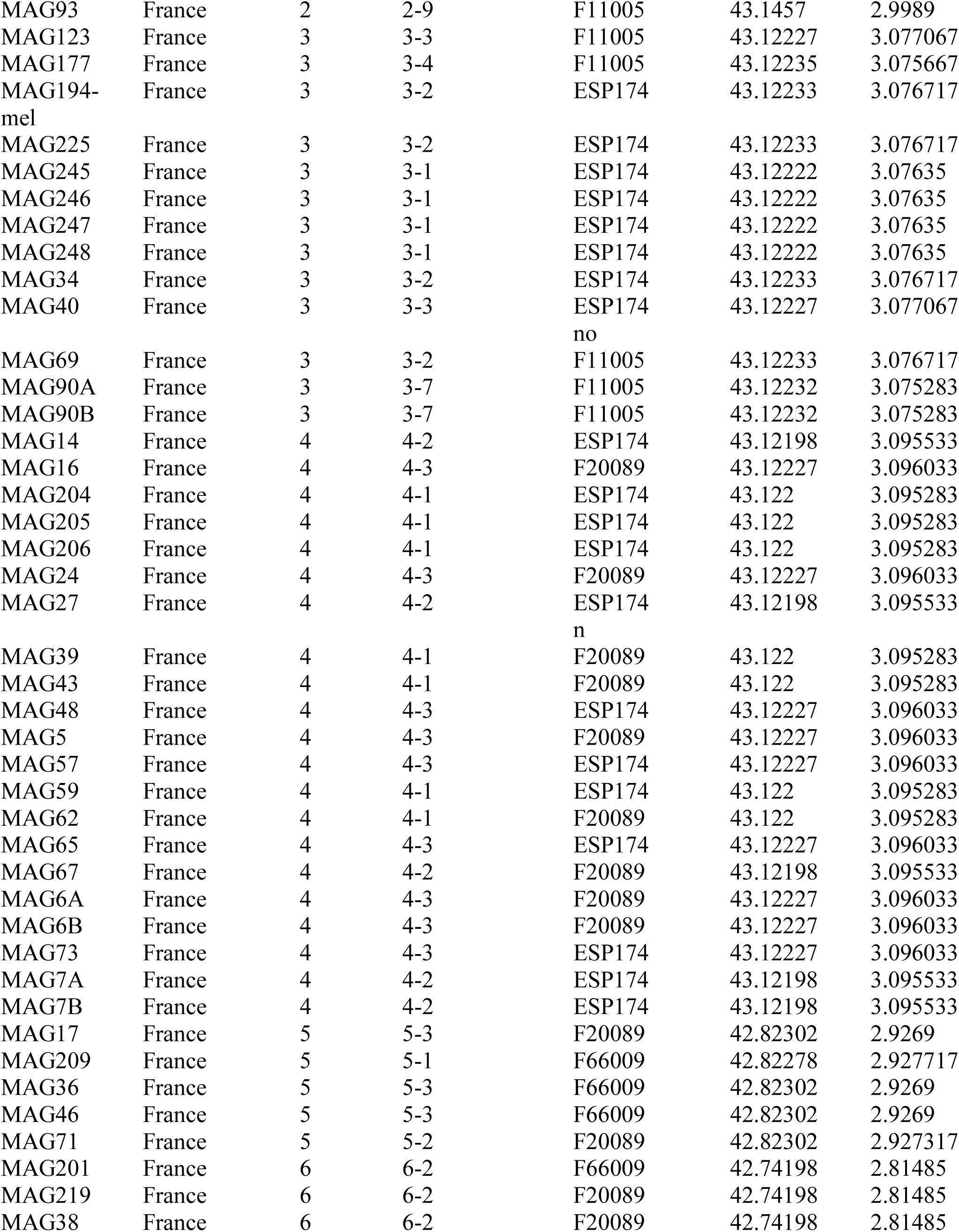
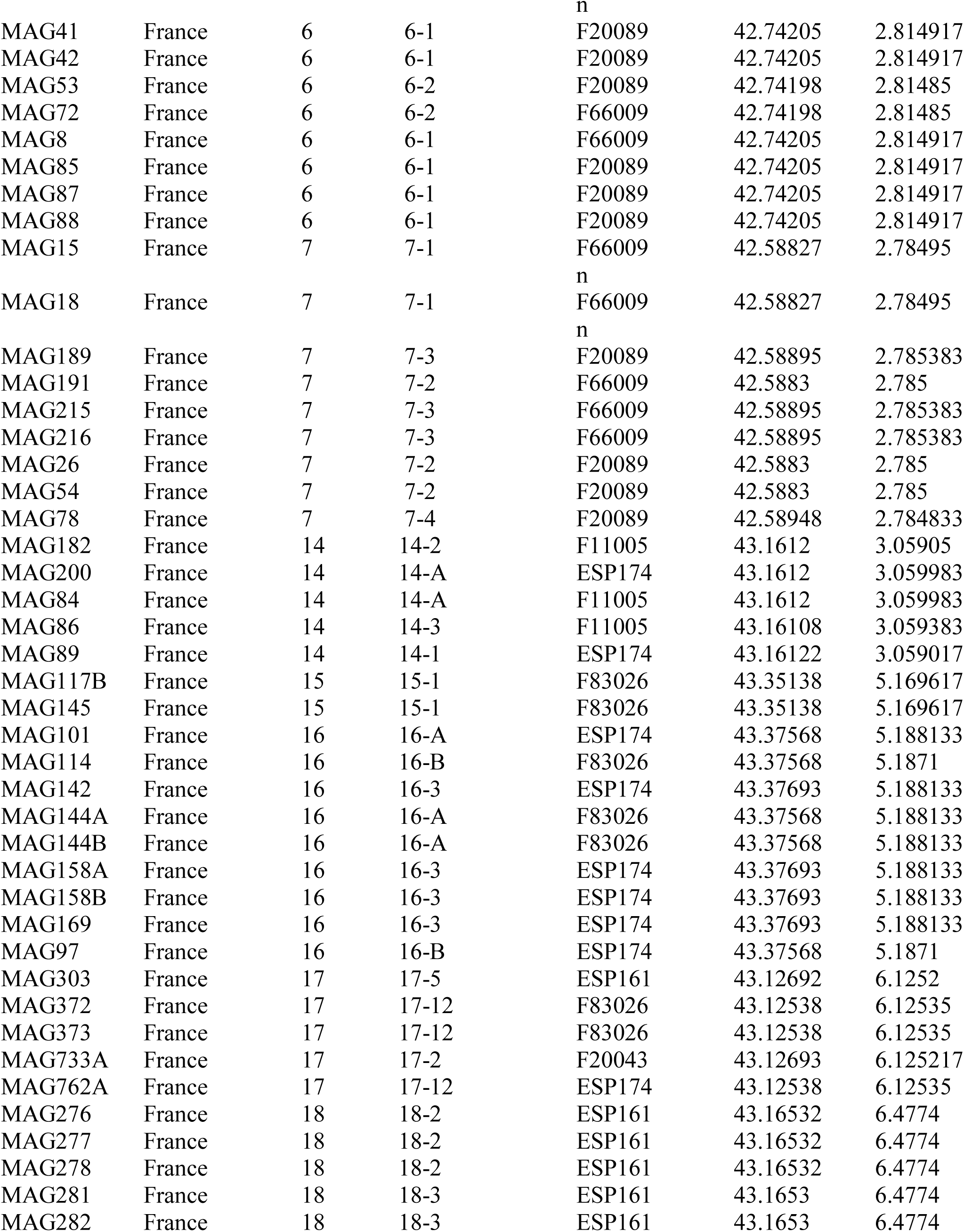
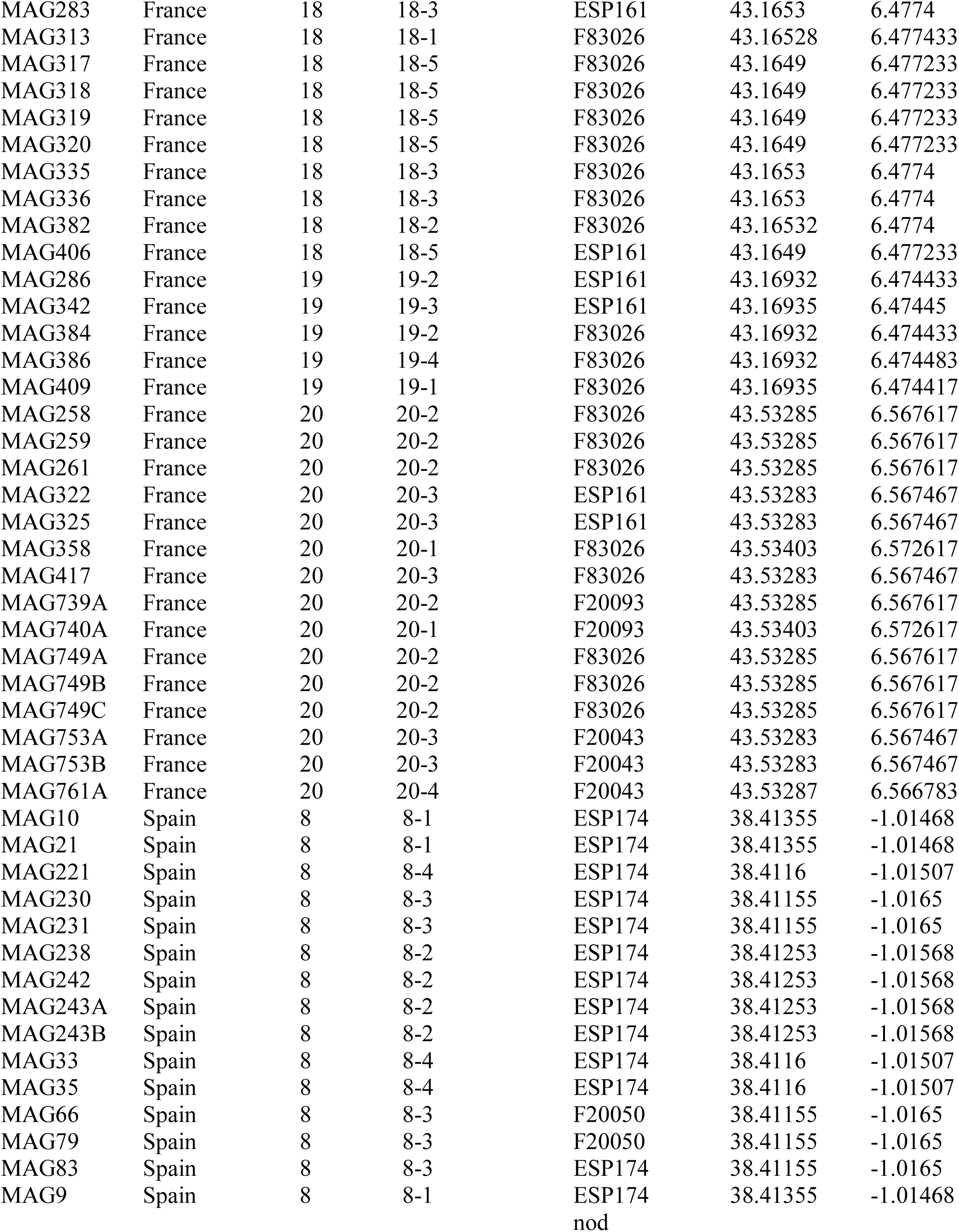
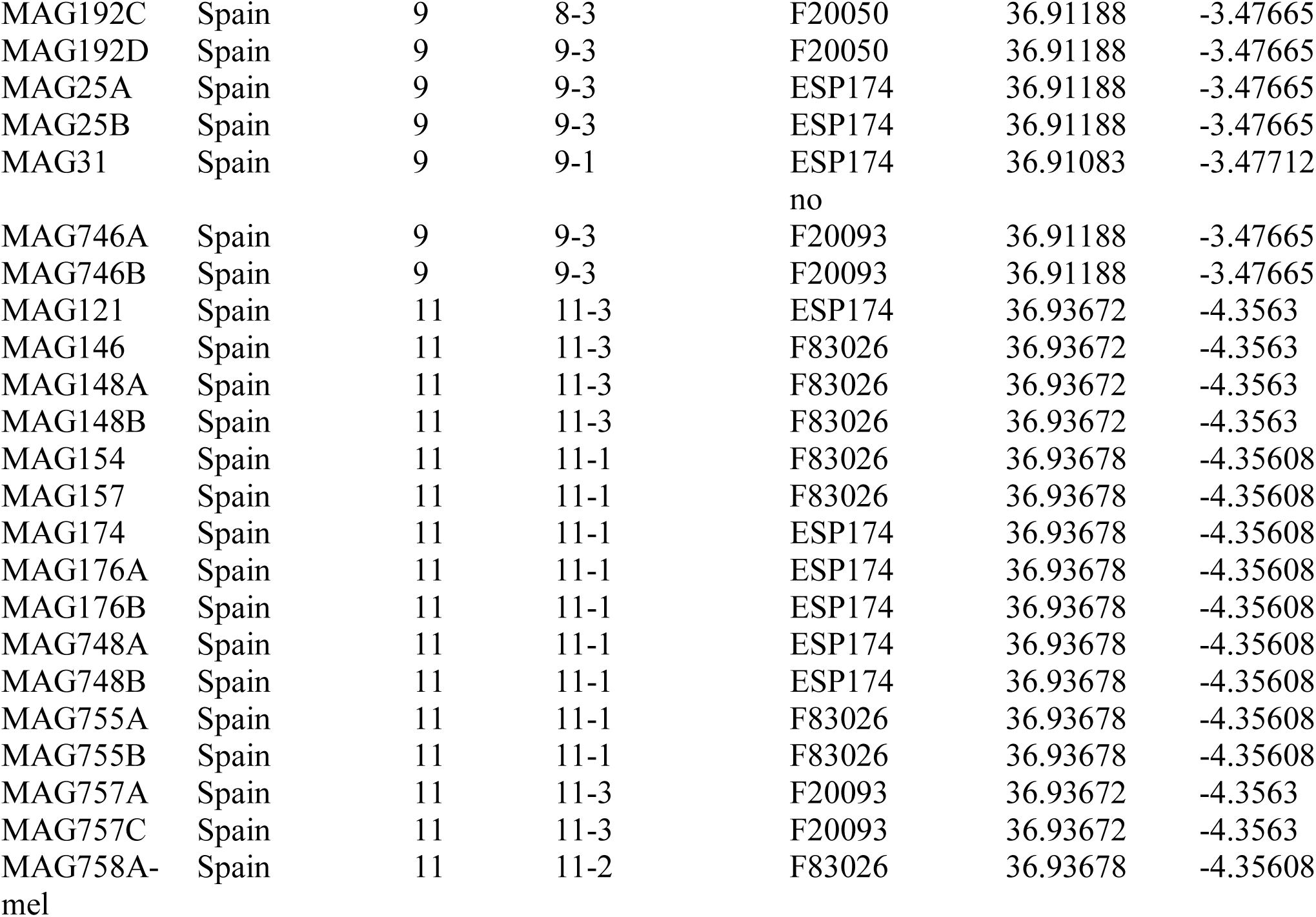
Metadata on strains of *Ensifer meliloti*. For each strain the region, sampling site (soil), and specific soil samples are given, along with the genotype of the *Medicago truncatula* plant used to isolate the strain, and the latitude and longitude at which the soil sample was taken.

**Table S2.** SNP counts for each element of the symbiont genome before any variant filtering, and after filtering for quality and depth using VCFtools.

|  | Chromosome | pSymA | pSymB |
| --- | --- | --- | --- |
| Pre-filtering | 414,004 | 179,505 | 222,935 |
| Quality filters | 34,689 | 15,162 | 22,460 |

**Table S3.** Matrices of population pairwise D_XY_ values for each *E. meliloti* genomic element. The populations are arranged from west to east (Spain-mainland France-Corsica). Cells background coloration indicates the level of D_XY_ (red = low values, green = high values).

*\*\*See supplementary Excel file*

## Supplemental figures

**Figure S1.**
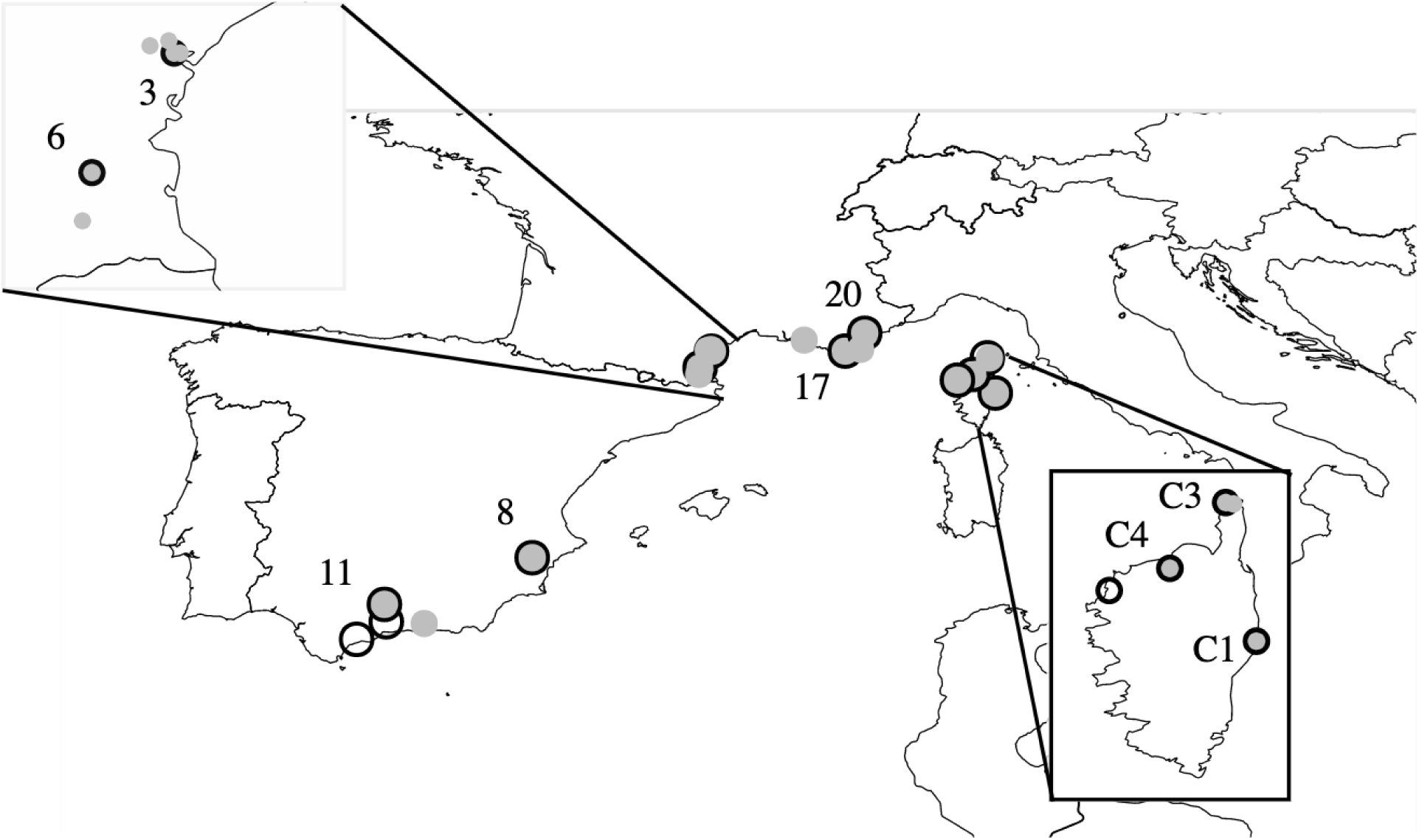
Map of 25 sampling locations in Spain, mainland France, and Corsica. Grey points with black outlines represent sites where both host and symbiont were sampled (n = 8), grey points represent sites where only symbionts were sampled (n = 13), and black circles represent sites where only hosts were sampled (n = 4).

**Figure S2.**
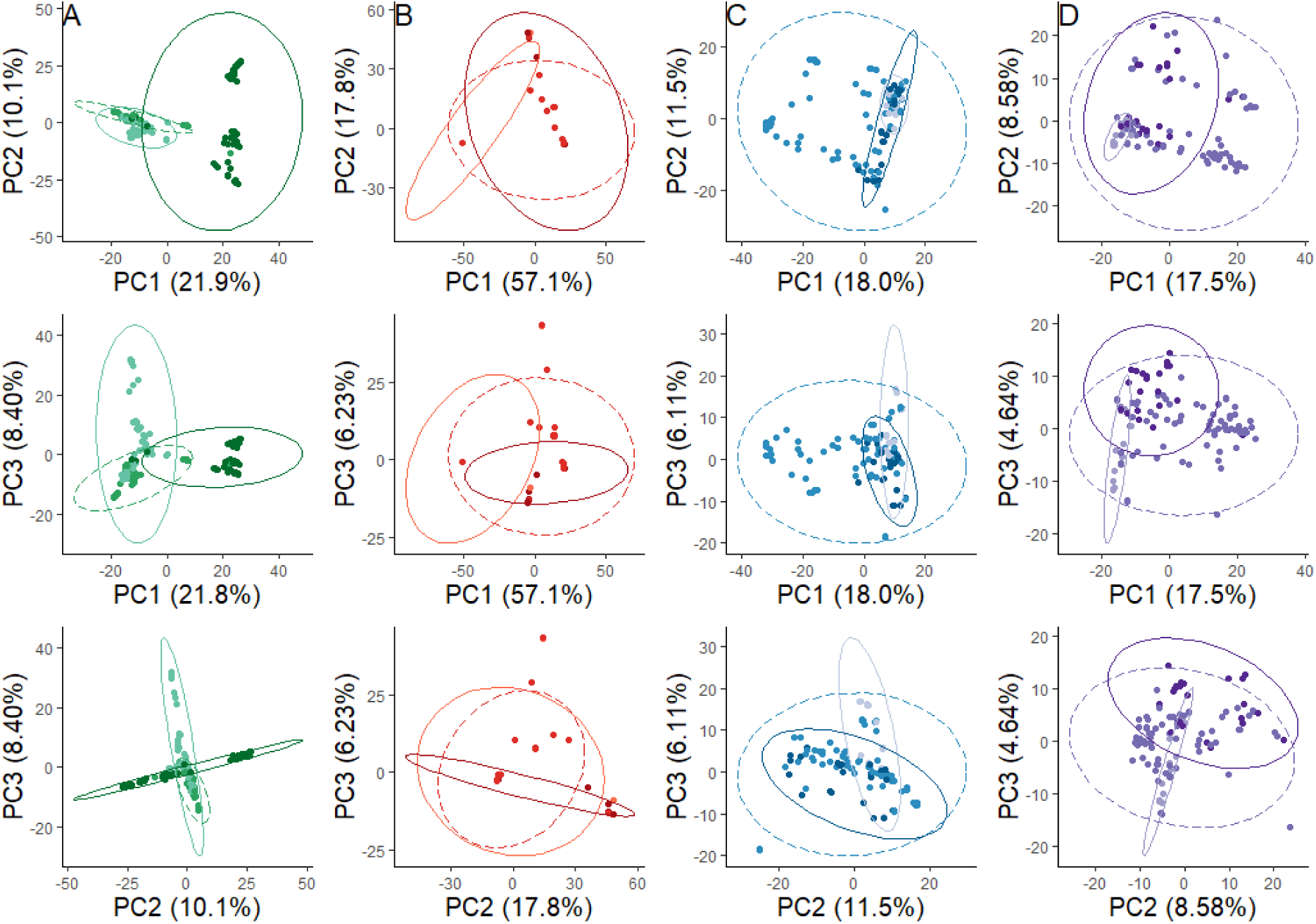
Principal component axis plots showing genome wide similarity on PCs 1-3 for the *Medicago* host plants (n=192) (**A**) and for symbiont (n=191) chromosome (**B**), pSymA (**C**), and pSymB (**D**). The darkest ellipse and points on each plot represent individuals from Spain, the intermediate color dashed line represent individuals from mainland France, and the lightest color represents individuals from Corsica.

**Figure S3.**
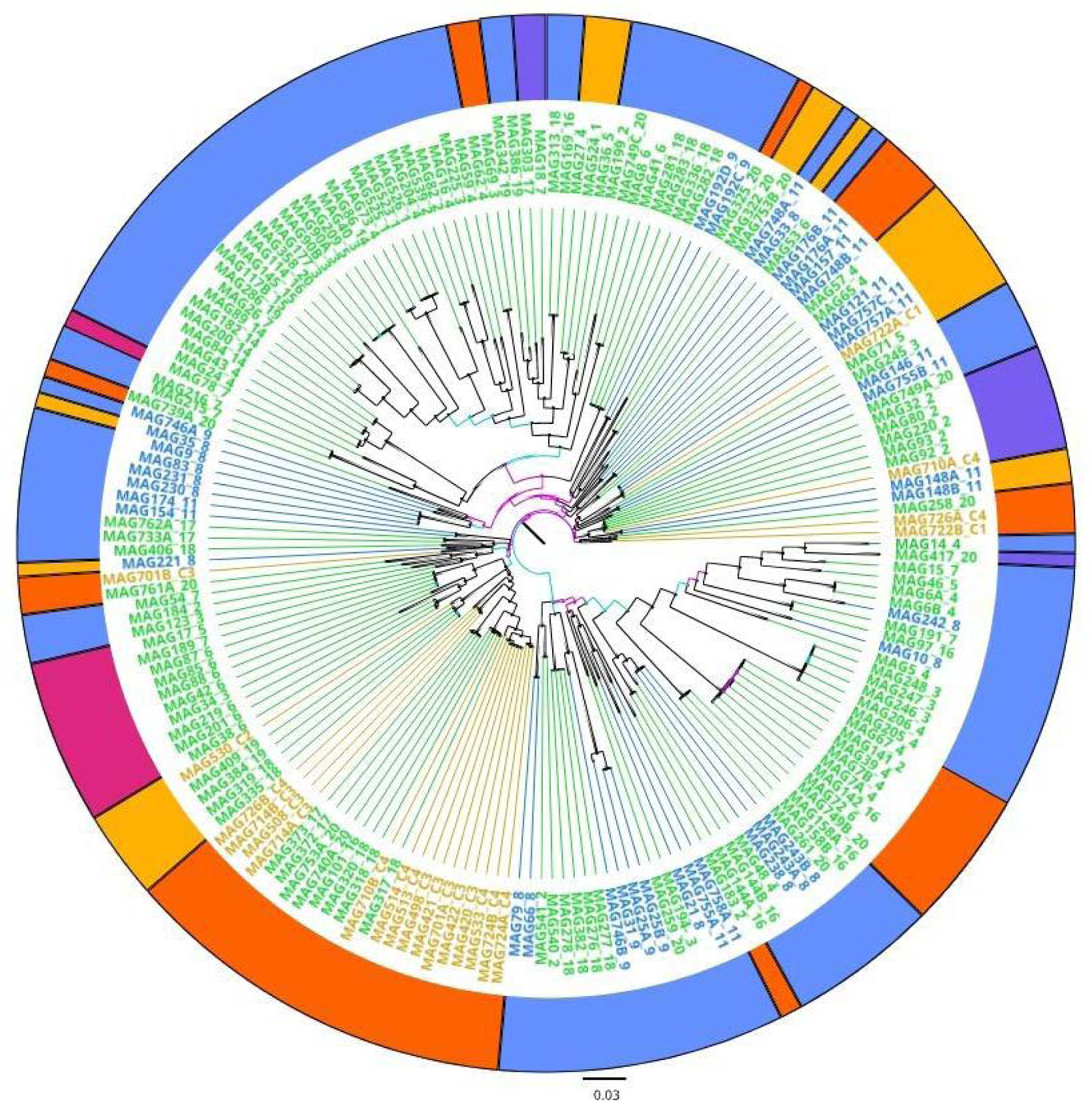
Neighbor joining tree of rhizobium individuals based on pSymB variant data. Individual tip labels (strain ID followed by soil population) are colored based on region of origin (orange from Corsica, green from mainland France, blue from Spain). Branch support indicated in teal (<70) or pink (<50). Outer ring represents the chromosomal cluster (from Fig. 2a) for comparison.

**Figure S4.**
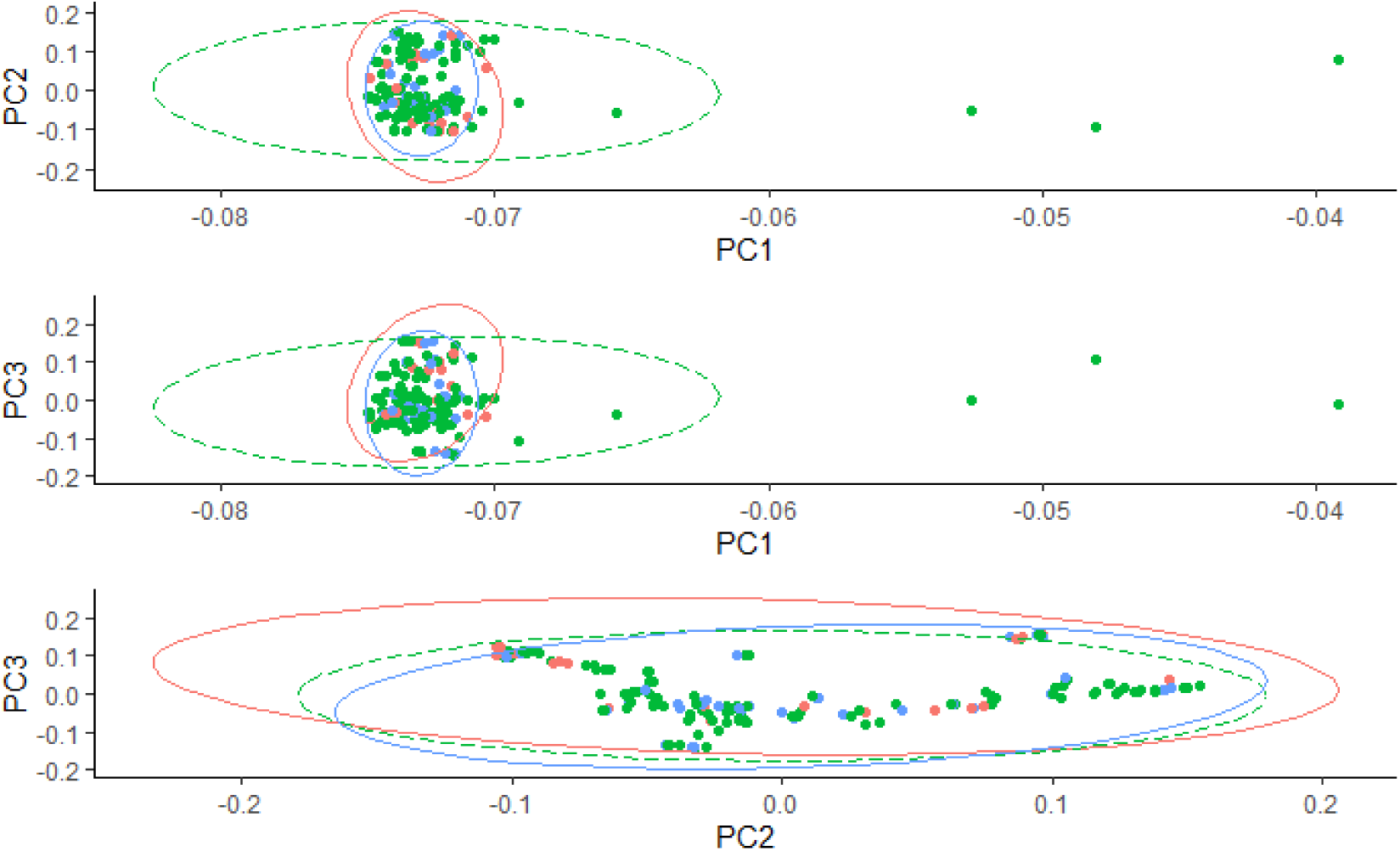
Principal component axis plot for all individuals of the symbiont (n=191) based on variable gene content. From top to bottom principal component 1 and 2, 1 and 3, and 2 and 3. The red ellipse and points on each plot represent individuals from Spain, the green dotted line and points represent individuals from mainland France, and the blue points and line represent individuals from Corsica.

**Figure S5.**
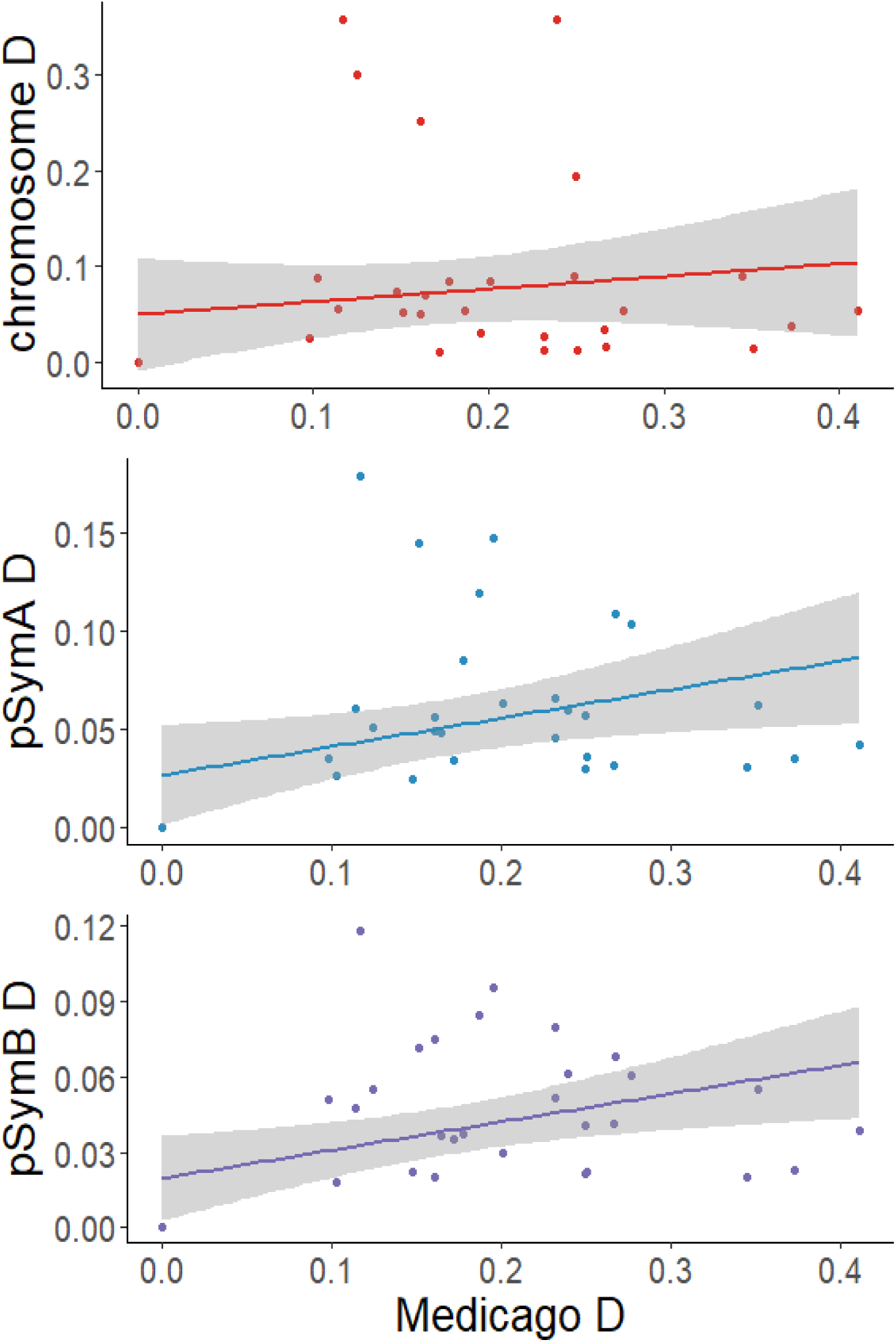
Tests of correlated population structure between the genomic elements in *E. meliloti* and the host plant *M. truncatula*

## References

Akçay, E. (2017). Population structure reduces benefits from partner choice in mutualistic symbiosis. Proceedings of the Royal Society B: Biological Sciences, 284(1850). doi: 10.1098/rspb.2016.2317

Anderson, B., Olivieri, I., Lourmas, M., & Stewart, B. A. (2004). Comparative population genetic structures and local adaptation of two mutualists. Evolution, 58(8), 1730–1747. doi: 10.1111/j.0014-3820.2004.tb00457.x

Bailly, X., Giuntini, E., Sexton, M. C., Lower, R. P. J., Harrison, P. W., Kumar, N., & Young, J. P. W. (2011). Population genomics of Sinorhizobium medicae based on low-coverage sequencing of sympatric isolates. ISME Journal, 5(11), 1722–1734. doi: 10.1038/ismej.2011.55

Bailly, X., Olivieri, I., De Mita, S., Cleyet-Marel, J. C., & Béna, G. (2006). Recombination and selection shape the molecular diversity pattern of nitrogen-fixing Sinorhizobium sp. associated to Medicago. Molecular Ecology, 15(10), 2719–2734. doi: 10.1111/j.1365-294X.2006.02969.x

Bankevich, A., Nurk, S., Antipov, D., Gurevich, A. A., Dvorkin, M., Kulikov, A. S., … Pevzner, P. A. (2012). SPAdes: A new genome assembly algorithm and its applications to single-cell sequencing. Journal of Computational Biology, 19(5), 455–477. doi: 10.1089/cmb.2012.0021

Batstone, R. T., Dutton, E. M., Wang, D., Yang, M., & Frederickson, M. E. (2017). The evolution of symbiont preference traits in the model legume Medicago truncatula. New Phytologist, 213(4), 1850–1861. doi: 10.1111/nph.14308

Batstone, R. T., O’Brien, A. M., Harrison, T. L., & Frederickson, M. E. (2020). Experimental evolution makes microbes more cooperative with their local host genotype. Science, 370(6515), 23–26. doi: 10.1126/science.abb7222

Baums, I. B., Devlin-Durante, M. K., & Lajeunesse, T. C. (2014). New insights into the dynamics between reef corals and their associated dinoflagellate endosymbionts from population genetic studies. Molecular Ecology, 23(17), 4203–4215. doi: 10.1111/mec.12788

Becker, A., Barnett, M. J., Capela, D., Dondrup, M., Kamp, P. B., Krol, E., … Goesmann, A. (2009). A portal for rhizobial genomes: RhizoGATE integrates a Sinorhizobium meliloti genome annotation update with postgenome data. Journal of Biotechnology, 140(1–2), 45–50. doi: 10.1016/j.jbiotec.2008.11.006

Berg, G. (2009). Plant-microbe interactions promoting plant growth and health: Perspectives for controlled use of microorganisms in agriculture. Applied Microbiology and Biotechnology, 84(1), 11–18. doi: 10.1007/s00253-009-2092-7

Biondi, E. G., Pilli, E., Giuntini, E., Roumiantseva, M. L., Andronov, E. E., Onichtchouk, O. P., … Bazzicalupo, M. (2003). Genetic relationship of Sinorhizobium meliloti and Sinorhizobium medicae strains isolated from Caucasian region. FEMS Microbiology Letters, 220(2), 207–213. doi: 10.1016/S0378-1097(03)00098-3

Blanca-Ordóñez, H., Oliva-García, J. J., Pérez-Mendoza, D., Soto, M. J., Olivares, J., Sanjuán, J., & Nogales, J. (2010). pSymA-dependent mobilization of the Sinorhizobium meliloti pSymB megaplasmid. Journal of Bacteriology, 192(23), 6309–6312. doi: 10.1128/JB.00549-10

Bonhomme, M., Boitard, S., Clemente, H. S., Dumas, B., Young, N., & Jacquet, C. (2015). Genomic signature of selective sweeps illuminates adaptation of medicago truncatula to root-associated microorganisms. Molecular Biology and Evolution, 32(8), 2097–2110. doi: 10.1093/molbev/msv092

Bonnin, I., Huguet, T., Gherardi, M., Prosperi, J.-M., & Olivieri, I. (1996). High level of polymorphism and spatial structure in a selfing plant species, Medicago truncatula (Leguminosae), shown using RAPD markers . American Journal of Botany, 83(7), 843–855. doi: 10.1002/j.1537-2197.1996.tb12776.x

Branca, A., Paape, T. D., Zhou, P., Briskine, R., Farmer, A. D., Mudge, J., … Tiffin, P. (2011). Whole-genome nucleotide diversity, recombination, and linkage disequilibrium in the model legume Medicago truncatula. Proceedings of the National Academy of Sciences of the United States of America, 108(42). doi: 10.1073/pnas.1104032108

Brown, S. P., Grillo, M. A., Podowski, J. C., & Heath, K. D. (2020). Soil origin and plant genotype structure distinct microbiome compartments in the model legume Medicago truncatula. Microbiome, 8(1), 1–17. doi: 10.1186/s40168-020-00915-9

Burghardt, L. T. (2020). Evolving together, evolving apart: measuring the fitness of rhizobial bacteria in and out of symbiosis with leguminous plants. New Phytologist, 228(1), 28–34. doi: 10.1111/nph.16045

Burghardt, L. T., Epstein, B., Guhlin, J., Nelson, M. S., Taylor, M. R., Young, N. D., … Tiffin, P. (2018). Select and resequence reveals relative fitness of bacteria in symbiotic and free-living environments. Proceedings of the National Academy of Sciences of the United States of America, 115(10), 2425–2430. doi: 10.1073/pnas.1714246115

Burghardt, L. T., Epstein, B., & Tiffin, P. (2019). Legacy of prior host and soil selection on rhizobial fitness in planta. Evolution, 73(9), 2013–2023. doi: 10.1111/evo.13807

Caldera, E. J., & Currie, C. R. (2012). The population structure of antibiotic-producing bacterial symbionts of apterostigma dentigerum ants: Impacts of coevolution and multipartite symbiosis. American Naturalist, 180(5), 604–617. doi: 10.1086/667886

Carlsson-Granér, U., & Thrall, P. H. (2015). Host resistance and pathogen infectivity in host populations with varying connectivity. Evolution, 69(4), 926–938. doi: 10.1111/evo.12631

Catchen, J., Hohenlohe, P. A., Bassham, S., Amores, A., & Cresko, W. A. (2013). StackCatchen, J., Hohenlohe, P. A., Bassham, S., Amores, A., & Cresko, W. A. (2013). Stacks: an analysis tool set for population genomics. Molecular Ecology, 22(11), 3124–3140. doi: 10.1111/mec.12354.Stacks

Cavassim, M. I. A., Moeskjær, S., Moslemi, C., Fields, B., Bachmann, A., Vilhjálmsson, B. J., … Andersen, S. U. (2020). Symbiosis genes show a unique pattern of introgression and selection within a rhizobium leguminosarum species complex. Microbial Genomics, 6(4). doi: 10.1099/mgen.0.000351

Chase, A., Arevalo, P., Brodie, E., Polz, M., Karaoz, U., & Martiny, J. B. H. (2019). Sympatric and allopatric differentiation delineates population structure in free-living terrestrial bacteria. MBio. doi: 10.1101/644468

Choudoir, M. J., Barberán, A., Menninger, H. L., Dunn, R. R., & Fierer, N. (2018). Variation in range size and dispersal capabilities of microbial taxa. Ecology, 99(2), 322–334. doi: 10.1002/ecy.2094

Cooper, V. S., Vohr, S. H., Wrocklage, S. C., & Hatcher, P. J. (2010). Why genes evolve faster on secondary chromosomes in bacteria. PLoS Computational Biology, 6(4). doi: 10.1371/journal.pcbi.1000732

Danecek, P., Auton, A., Abecasis, G., Albers, C. A., Banks, E., DePristo, M. A., … Durbin, R. (2011). The variant call format and VCFtools. Bioinformatics, 27(15), 2156–2158. doi: 10.1093/bioinformatics/btr330

Degnan, J. H., & Rosenberg, N. A. (2009). Gene tree discordance, phylogenetic inference and the multispecies coalescent. Trends in Ecology and Evolution, 24(6), 332–340. doi: 10.1016/j.tree.2009.01.009

diCenzo, G. C., & Finan, T. M. (2017). The Divided Bacterial Genome. Microbiology and Molecular Biology Reviews, 81(3), 1–37.

diCenzo, G. C., MacLean, A. M., Milunovic, B., Golding, G. B., & Finan, T. M. (2014). Examination of Prokaryotic Multipartite Genome Evolution through Experimental Genome Reduction. PLoS Genetics, 10(10). doi: 10.1371/journal.pgen.1004742

Ding, H., & Hynes, M. F. (2009). Plasmid transfer systems in the rhizobia. Canadian Journal of Microbiology, 55(8), 917–927. doi: 10.1139/W09-056

Dray, S., & Dufour, A. B. (2007). The ade4 package: Implementing the duality diagram for ecologists. Journal of Statistical Software, 22(4), 1–20. doi: 10.18637/jss.v022.i04

Dybdahl, M. F., & Lively, C. M. (1996). The geography of coevolution: Comparative population structures for a snail and its trematode parasite. Evolution, 50(6), 2264–2275. doi: 10.1111/j.1558-5646.1996.tb03615.x

Epstein, B., Branca, A., Mudge, J., Bharti, A. K., Briskine, R., Farmer, A. D., … Tiffin, P. (2012). Population Genomics of the Facultatively Mutualistic Bacteria Sinorhizobium meliloti and S. medicae. PLoS Genetics, 8(8), 1–10. doi: 10.1371/journal.pgen.1002868

Epstein, B., Sadowsky, M. J., & Tiffin, P. (2014). Selection on Horizontally Transferred and Duplicated Genes in Sinorhizobium (Ensifer), the Root-Nodule Symbionts of Medicago. Genome Biology and Evolution, 6(5), 1199–1209. doi: 10.1093/gbe/evu090

Epstein, B., & Tiffin, P. (2021). Comparative genomics reveals high rates of horizontal transfer and strong purifying selection on rhizobial symbiosis genes. Proceedings of the Royal Society B: Biological Sciences, 288(1942). doi: 10.1098/rspb.2020.1804

Fernandes, L. D., Lemos-Costa, P., Guimarães, P. R., Thompson, J. N., & de Aguiar, M. A. M. (2019). Coevolution Creates Complex Mosaics across Large Landscapes. The American Naturalist, 194(2), 000–000. doi: 10.1086/704157

Frederickson, M. E. (2017). Mutualisms Are Not on the Verge of Breakdown. Trends in Ecology and Evolution, 32(10), 727–734. doi: 10.1016/j.tree.2017.07.001

Friesen, M. L., & Mathias, A. (2010). Mixed infections may promote diversification of mutualistic symbionts: Why are there ineffective rhizobia? Journal of Evolutionary Biology, 23(2), 323–334. doi: 10.1111/j.1420-9101.2009.01902.x

Friesen, M.L. (2012). Widespread fitness alignment in the legume-rhizobium symbiosis. New Phytologist, 194(4), 1096–1111. doi: 10.1111/j.1469-8137.2012.04099.x

Galardini, M., Pini, F., Bazzicalupo, M., Biondi, E. G., & Mengoni, A. (2013). Replicon-dependent bacterial genome evolution: The case of Sinorhizobium meliloti. Genome Biology and Evolution, 5(3), 542–558. doi: 10.1093/gbe/evt027

Galibert, F., Finan, T. M., Long, S. R., Pühler, A., Abola, P., Ampe, F., … Batut, J. (2001). The composite genome of the legume symbiont Sinorhizobium meliloti. Science, 293(5530), 668–672. doi: 10.1126/science.1060966

Gandon, S., Capowiez, Y., Dubois, Y., Michalakis, Y., & Olivieri, I. (1996a). Local adaptation and gene-for-gene coevolution in a metapopulation model. Proceedings of the Royal Society B: Biological Sciences, 263(1373), 1003–1009. Retrieved from https://royalsocietypublishing.org/doi/10.1098/rspb.1996.0148

Gandon, S., Capowiez, Y., Dubois, Y., Michalakis, Y., & Olivieri, I. (1996b). Local adaptation and gene-for-gene coevolution in a metapopulation model. Proceedings of the Royal Society B: Biological Sciences, 263(1373), 1003–1009. doi: https://doi.org/10.1098/rspb.1996.0148

Gano-Cohen, K. A., Wendlandt, C. E., Moussawi, K. Al, Stokes, P. J., Quides, K. W., Weisberg, A. J., … Sachs, J. L. (2020). Recurrent mutualism breakdown events in a legume rhizobia metapopulation. Proceedings of the Royal Society B: Biological Sciences, 287(1919). doi: 10.1098/rspb.2019.2549

Garrison, E., & Marth, G. (2012). Haplotype-based variant detection from short-read sequencing. arXiv preprint, http://arxiv.org/abs/1207.3907

Green, J. L., Bohannan, B. J. M., & Whitaker, R. J. (2008). Microbial biogeography: From taxonomy to traits. Science, 320(5879), 1039–1043. doi: 10.1126/science.1153475

Grillo, M. A., De Mita, S., Burke, P. V., Solórzano-Lowell, K. L. S., & Heath, K. D. (2016). Intrapopulation genomics in a model mutualist: Population structure and candidate symbiosis genes under selection in *Medicago truncatula*. Evolution, 70(12), 2704–2717. doi: 10.1111/evo.13095

Hanson, C. A., Fuhrman, J. A., Horner-Devine, M. C., & Martiny, J. B. H. (2012). Beyond biogeographic patterns: Processes shaping the microbial landscape. Nature Reviews Microbiology, 10(7), 497–506. doi: 10.1038/nrmicro2795

Harrison, P. W., Lower, R. P. J., Kim, N. K. D., & Young, J. P. W. (2010). Introducing the bacterial ‘chromid’: not a chromosome, not a plasmid. Trends in Microbiology, 18(4), 141–148. doi: 10.1016/J.TIM.2009.12.010

Harrison, T. L., Wood, C. W., Heath, K. D., & Stinchcombe, J. R. (2017a). Geographically structured genetic variation in the Medicago lupulina–Ensifer mutualism. Evolution, 71(7), 1787–1801. doi: 10.1111/evo.13268

Harrison, T. L., Wood, C. W., Heath, K. D., & Stinchcombe, J. R. (2017b). Geographically structured genetic variation in the *Medicago lupulina* - *Ensifer* mutualism. Evolution, 71(7), 1787–1801. doi: 10.1111/evo.13268

Heath, K. D. (2010). Intergenomic epistasis and coevolutionary constraint in plants and rhizobia. Evolution, 64(5), 1446–1458. doi: 10.1111/j.1558-5646.2009.00913.x

Heath, K. D., & Grillo, M. A. (2016). Rhizobia: tractable models for bacterial evolutionary ecology. Environmental Microbiology, 18(12), 4307–4311. doi: 10.1111/1462-2920.13492

Heath, K. D., & Stinchcombe, J. R. (2014a). Explaining mutualism variation: A new evolutionary paradox? Evolution, 68(2), 309–317. doi: 10.1111/evo.12292

Heath, K. D., & Tiffin, P. (2009). Stabilizing mechanisms in a legume-rhizobium mutualism. Evolution, 63(3), 652–662. doi: 10.1111/j.1558-5646.2008.00582.x

Hoetzinger, M., Pitt, A., Huemer, A., & Hahn, M. W. (2021). Continental-Scale Gene Flow Prevents Allopatric Divergence of Pelagic Freshwater Bacteria. Genome Biology and Evolution, 13(3), 1–18. doi: 10.1093/gbe/evab019

Hollowell, A. C., Regus, J. U., Gano, K. A., Bantay, R., Centeno, D., Pham, J., … Sachs, J. L. (2016). Epidemic Spread of Symbiotic and Non-Symbiotic Bradyrhizobium Genotypes Across California. Microbial Ecology, 71(3), 700–710. doi: 10.1007/s00248-015-0685-5

Hollowell, Amanda C., Regus, J. U., Turissini, D., Gano-Cohen, K. A., Bantay, R., Bernardo, A., … Sachs, J. L. (2016). Metapopulation dominance and genomicisland acquisition of Bradyrhizobium with superior catabolic capabilities. Proceedings of the Royal Society B: Biological Sciences, 283(1829). doi: 10.1098/rspb.2016.0496

Jombart, T., & Ahmed, I. (2011). adegenet 1.3-1: New tools for the analysis of genome-wide SNP data. Bioinformatics, 27(21), 3070–3071. doi: 10.1093/bioinformatics/btr521

Jones, E. I., Afkhami, M. E., Akçay, E., Bronstein, J. L., Bshary, R., Frederickson, M. E., … Friesen, M. L. (2015). Cheaters must prosper: Reconciling theoretical and empirical perspectives on cheating in mutualism. Ecology Letters, 18(11), 1270–1284. doi: 10.1111/ele.12507

Kamvar, Z. N., Tabima, J. F., & Gr unwald, N. J. (2014). Poppr: An R package for genetic analysis of populations with clonal, partially clonal, and/or sexual reproduction. PeerJ, 2014(1), 1–14. doi: 10.7717/peerj.281

Kaneko, T., Nakamura, Y., Sato, S., Minamisawa, K., Uchiumi, T., Sasamoto, S., … Tabata, S. (2002). Complete genomic sequence of nitrogen-fixing symbiotic bacterium Bradyrhizobium japonicum USDA110. DNA Research, 9(6), 189–197. doi: 10.1093/dnares/9.6.189

Klinger, C. R., Lau, J. A., & Heath, K. D. (2016). Ecological genomics of mutualism decline in nitrogen-fixing bacteria. Proceedings of the Royal Society B: Biological Sciences, 283(1826), 20152563. doi: 10.1098/rspb.2015.2563

Koppell, J. H., & Parker, M. A. (2012). Phylogenetic clustering of Bradyrhizobium symbionts on legumes indigenous to North America. Microbiology (United Kingdom), 158(8), 2050–2059. doi: 10.1099/mic.0.059238-0

Kumar, N., Lad, G., Giuntini, E., Kaye, M. E., Udomwong, P., Shamsani, N. J., … Bailly, X. (2015). Bacterial genospecies that are not ecologically coherent: population genomics of Rhizobium leguminosarum. Open Biology, 5(1), 140133–140133. doi: 10.1098/rsob.140133

Laine, A. L. (2005). Spatial scale of local adaptation in a plant-pathogen metapopulation. Journal of Evolutionary Biology, 18(4), 930–938. doi: 10.1111/j.1420-9101.2005.00933.x

Li, H., & Durbin, R. (2009). Fast and accurate short read alignment with Burrows-Wheeler transform. Bioinformatics, 25(14), 1754–1760. doi: 10.1093/bioinformatics/btp324

Locey, K. J., & Lennon, J. T. (2016). Scaling laws predict global microbial diversity. Proceedings of the National Academy of Sciences of the United States of America, 113(21), 5970–5975. doi: 10.1073/pnas.1521291113

Martiny, J. B. H., Bohannan, B. J. M., Brown, J. H., Colwell, R. K., Fuhrman, J. A., Green, J. L., … Staley, J. T. (2006). Microbial biogeography: Putting microorganisms on the map. Nature Reviews Microbiology, 4(2), 102–112. doi: 10.1038/nrmicro1341

Masatoshi, N. (1972). Genetic Distance between Populations. American Naturalist, 106(949), 283–292.

Nelson, M., Guhlin, J., Epstein, B., Tiffin, P., & Sadowsky, M. J. (2018a). The complete replicons of 16 Ensifer meliloti strains offer insights into intra- and inter-replicon gene transfer, transposon-associated loci, and repeat elements. Microbial Genomics, 4(5). doi: 10.1099/mgen.0.000174

Nelson, M., Guhlin, J., Epstein, B., Tiffin, P., & Sadowsky, M. J. (2018b). The complete replicons of 16 Ensifer meliloti strains offer insights into intra- and inter-replicon gene transfer, transposon-associated loci, and repeat elements. Microbial Genomics, 4(5), 1–11. doi: 10.1099/mgen.0.000174

Nuismer, S. L., Thompson, J. N., & Gomulkiewicz, R. (2003). Coevolution between hosts and parasites with partially overlapping geographic ranges. Journal of Evolutionary Biology, 16(6), 1337–1345. doi: 10.1046/j.1420-9101.2003.00609.x

Ossler, J. N., & Heath, K. D. (2018). Shared genes but not shared genetic variation: Legume colonization by two belowground symbionts. American Naturalist, 191(3), 395–406. doi: 10.1086/695829

Page, A. J., Cummins, C. A., Hunt, M., Wong, V. K., Reuter, S., Holden, M. T. G., … Parkhill, J. (2015). Roary: Rapid large-scale prokaryote pan genome analysis. Bioinformatics, 31(22), 3691–3693. doi: 10.1093/bioinformatics/btv421

Paradis, E., & Schliep, K. (2019). Ape 5.0: An environment for modern phylogenetics and evolutionary analyses in R. Bioinformatics, 35(3), 526–528. doi: 10.1093/bioinformatics/bty633

Parker, M. A. (1999). Mutualism in metapopulations of legumes and rhizobia. American Naturalist, 153(SUPPL.), 48–60. doi: 10.1086/303211

Pembleton, L. W., Cogan, N. O. I., & Forster, J. W. (2013). StAMPP: An R package for calculation of genetic differentiation and structure of mixed-ploidy level populations. Molecular Ecology Resources, 13(5), 946–952. doi: 10.1111/1755-0998.12129

Pérez-Mendoza, D., Sepúlveda, E., Pando, V., Muñoz, S., Nogales, J., Olivares, J., … Sanjuán, J. (2005). Identification of the rctA gene, which is required for repression of conjugative transfer of rhizobial symbiotic megaplasmids. Journal of Bacteriology, 187(21), 7341–7350. doi: 10.1128/JB.187.21.7341-7350.2005

Pérez Carrascal, O. M., VanInsberghe, D., Juárez, S., Polz, M. F., Vinuesa, P., & González, V. (2016). Population genomics of the symbiotic plasmids of sympatric nitrogen-fixing *Rhizobium* species associated with *Phaseolus vulgaris*. Environmental Microbiology, 18(8), 2660–2676. doi: 10.1111/1462-2920.13415

Pita, L., Rix, L., Slaby, B. M., Franke, A., & Hentschel, U. (2018). The sponge holobiont in a changing ocean: from microbes to ecosystems. Microbiome, 6(1), 46. doi: 10.1186/s40168-018-0428-1

Porter, S. S., Faber-Hammond, J., Montoya, A. P., Friesen, M. L., & Sackos, C. (2019). Dynamic genomic architecture of mutualistic cooperation in a wild population of Mesorhizobium. ISME Journal, 13(2), 301–315. doi: 10.1038/s41396-018-0266-y

Porter, S. S., & Simms, E. L. (2014). Selection for cheating across disparate environments in the legume-rhizobium mutualism. Ecology Letters, 17(9), 1121–1129. doi: 10.1111/ele.12318

Pretorius-Guth, I. M., Puhler, A., & Simon, R. (1990). Conjugal transfer of megaplasmid 2 between Rhizobium meliloti strains in alfalfa nodules. Applied and Environmental Microbiology, 56(8), 2354–2359. doi: 10.1128/aem.56.8.2354-2359.1990

Rambaut, A. (2018). FigTree. Edinburgh. Retrieved from http://tree.bio.ed.ac.uk/software/figtree/

Revillini, D., Gehring, C. A., & Johnson, N. C. (2016). The role of locally adapted mycorrhizas and rhizobacteria in plant–soil feedback systems. Functional Ecology, 30(7), 1086–1098. doi: 10.1111/1365-2435.12668

Reynolds, H. L., Packer, A., Bever, J. D., & Clay, K. (2003). Grassroots ecology: Plant-microbe-soil interactions as drivers of plant community structure and dynamics. Ecology, 84(9), 2281–2291. doi: 10.1890/02-0298

[dataset] Riley, A.B., Grillo, M.A., Epstein, B., Tiffin, P., & Heath, K.D. (2021). Partners in space: Discordant population structure between legume hosts and rhizobium symbionts in their native range. NCBI - and DRYAD #.

Ronfort, J., Bataillon, T., Santoni, S., Delalande, M., David, J. L., & Prosperi, J. M. (2006). Microsatellite diversity and broad scale geographic structure in a model legume: Building a set of nested core collection for studying naturally occurring variation in Medicago truncatula. BMC Plant Biology, 6, 1–13. doi: 10.1186/1471-2229-6-28

Rudman, S. M., Barbour, M. A., Csilléry, K., Gienapp, P., Guillaume, F., Hairston, N. G., … Levine, J. M. (2018). What genomic data can reveal about eco-evolutionary dynamics. Nature Ecology and Evolution, 2(1), 9–15. doi: 10.1038/s41559-017-0385-2

Sachs, J. L., Quides, K. W., & Wendlandt, C. E. (2018). Legumes versus rhizobia□ : a model for ongoing conflict in symbiosis. New Phytologist, 219, 1199–1206. doi: 10.1111/nph.15222

Savolainen, O., Lascoux, M., & Merilä, J. (2013). Ecological genomics of local adaptation. Nature Reviews Genetics, 14(11), 807–820. doi: 10.1038/nrg3522

Seemann, T. (2014). Prokka: Rapid prokaryotic genome annotation. Bioinformatics, 30(14), 2068–2069. doi: 10.1093/bioinformatics/btu153

Siol, M., Prosperi, J. M., Bonnin, I., & Ronfort, J. (2008). How multilocus genotypic pattern helps to understand the history of selfing populations: A case study in Medicago truncatula. Heredity, 100(5), 517–525. doi: 10.1038/hdy.2008.5

Smith, C. I., Godsoe, W. K. W., Tank, S., Yoder, J. B., & Pellmyr, O. (2008). Distinguishing coevolution from covicariance in an obligate pollination mutualism: Asynchronous divergence in Joshua tree and its pollinators. Evolution, 62(10), 2676–2687. doi: 10.1111/j.1558-5646.2008.00500.x

Stoy, K. S., Gibson, A. K., Gerardo, N. M., & Morran, L. T. (2020). A need to consider the evolutionary genetics of host–symbiont mutualisms. Journal of Evolutionary Biology, 33(12), 1656–1668. doi: 10.1111/jeb.13715

Strobel, H. M., Alda, F., Sprehn, C. G., Blum, M. J., & Heins, D. C. (2016). Geographic and host-mediated population genetic structure in a cestode parasite of the three-spined stickleback. Biological Journal of the Linnean Society, 119(2), 381–396. doi: 10.1111/bij.12826

Tack, A. J. M., Horns, F., & Laine, A. L. (2014). The impact of spatial scale and habitat configuration on patterns of trait variation and local adaptation in a wild plant parasite. Evolution, 68(1), 176–189. doi: 10.1111/evo.12239

Tang, H., Krishnakumar, V., Bidwell, S., Rosen, B., Chan, A., Zhou, S., … Town, C. D. (2014). An improved genome release (version Mt4.0) for the model legume Medicago truncatula. BMC Genomics, 15(1), 1–14. doi: 10.1186/1471-2164-15-312

Thompson, A. R., Thacker, C. E., & Shaw, E. Y. (2005). Phylogeography of marine mutualists: Parallel patterns of genetic structure between obligate goby and shrimp partners. Molecular Ecology, 14(11), 3557–3572. doi: 10.1111/j.1365-294X.2005.02686.x

Thompson, J. N. (1994). The Coevolutionary Process (1st ed.). Chicago, IL, USA: The University Of Chicago Press Books.

Thompson, J. N. (2005). The Geographic Mosaic of Coevolution (1st ed.). Chicago Press.

Thrall, P. H., Hochberg, M. E., Burdon, J. J., & Bever, J. D. (2007). Coevolution of symbiotic mutualists and parasites in a community context. Trends in Ecology and Evolution, 22(3), 120–126. doi: 10.1016/j.tree.2006.11.007

VanInsberghe, D., Arevalo, P., Chien, D., & Polz, M. F. (2020). How can microbial population genomics inform community ecology? Philosophical Transactions of the Royal Society B: Biological Sciences, 375(1798). doi: 10.1098/rstb.2019.0253

Vincent, J. (1970). A Manual for the Practical Study of Root-Nodule Bacteria. Oxford and Edinburgh: Blackwell Scientific.

Vos, M., & Velicer, G. J. (2008). Isolation by Distance in the Spore-Forming Soil Bacterium Myxococcus xanthus. Current Biology, 18(5), 386–391. doi: 10.1016/J.CUB.2008.02.050

Whitaker, R. J., & Banfield, J. F. (2006). Population genomics in natural microbial communities. Trends in Ecology and Evolution, 21(9), 508–516. doi: 10.1016/j.tree.2006.07.001

Wickham, H. (2016). ggplot2: Elegant Graphics for Data Analysis. New York: Springer-Verlag. Retrieved from https://ggplot2.tidyverse.org

Wood, C. W., Pilkington, B. L., Vaidya, P., Biel, C., & Stinchcombe, J. R. (2018). Genetic conflict with a parasitic nematode disrupts the legume-rhizobia mutualism. Evolution Letters, 2(3), 233–245. doi: 10.1002/evl3.51

Yates, R., Howieson, J., De Meyer, S. E., Tian, R., Seshadri, R., Pati, A., … Reeve, W. (2015). High-quality permanent draft genome sequence of Rhizobium sullae strain WSM1592; a Hedysarum coronarium microsymbiont from Sassari, Italy. Standards in Genomic Sciences, 10(JULY2015), 1–6. doi: 10.1186/s40793-015-0020-2

Yoder, J. B., & Nuismer, S. L. (2010). When does coevolution promote diversification? American Naturalist, 176(6), 802–817. doi: 10.1086/657048

Yoder, J. B., Stanton-Geddes, J., Zhou, P., Briskine, R., Young, N. D., & Tiffin, P. (2014). Genomic signature of adaptation to climate in Medicago truncatula. Genetics, 196(4), 1263–1275. doi: 10.1534/genetics.113.159319

Young, J. P. W., Crossman, L. C., Johnston, A. W. B., Thomson, N. R., Ghazoui, Z. F., Hull, K. H., … Parkhill, J. (2006). The genome of Rhizobium leguminosarum has recognizable core and accessory components. Genome Biology, 7(4). doi: 10.1186/gb-2006-7-4-r34

Young, J. P. W., & Haukka, K. E. (1996). Diversity and phylogeny of rhizobia. New Phytologist, 133(1), 87–94. doi: 10.1111/j.1469-8137.1996.tb04344.x

Younginger, B. S., & Friesen, M. L. (2019). Connecting signals and benefits through partner choice in plant-microbe interactions. FEMS Microbiology Letters, 366(18), 1–11. doi: 10.1093/femsle/fnz217

Zribi, K., Mhamdi, R., Huguet, T., & Aouani, M. E. (2004). Distribution and genetic diversity of rhizobia nodulating natural populations of Medicago truncatula in tunisian soils. Soil Biology and Biochemistry, 36(6), 903–908. doi: 10.1016/j.soilbio.2004.02.003

